# A nuclear role for the Argonaute protein AGO2 in mammalian gametogenesis

**DOI:** 10.1101/2021.08.17.456253

**Authors:** Kimberly N Griffin, Haixin Li, Benjamin William Walters, Huafeng Wang, Carolyn B Kaya, Jean Kanyo, TuKiet Lam, Andy L Cox, Jean-Ju Chung, Bluma J Lesch

## Abstract

Argonaute 2 (AGO2) is a ubiquitously expressed protein critical for regulation of mRNA translation and vital to animal development. AGO2 protein is found in both cytoplasmic and nuclear compartments, and while its cytoplasmic role is well studied, the biological relevance of nuclear AGO2 is unclear. Here, we address this problem in vivo, using developing spermatogenic cells as a model. Remarkably, we find that AGO2 acts in the germ cell nucleus to positively regulate protein expression. We show that AGO2 dynamically binds both chromatin and nuclear mRNA transcripts of hundreds of genes required for sperm production, and germline conditional knockout (cKO) of *Ago2* causes depletion of the corresponding proteins, along with defects in sperm number and morphology. Nuclear AGO2 partners with splicing, export, and chromatin factors to promote transcript export and protein expression. Together, our data reveal an unexpected role for nuclear AGO2 in enhancing expression of developmentally important genes.

## Introduction

Cells control gene expression through multiple mechanisms, including regulation of chromatin state and transcription in the nucleus, mRNA export from the nucleus to the cytoplasm, and cytoplasmic translation from mRNA to protein. Argonaute (AGO) proteins play a major role in fine-tuning gene expression through posttranscriptional regulation (Ebert and Sharp, 2012; Hornstein and Shomron, 2006; Hutvágner and Zamore, 2002; Lee et al., 2004; Lim et al., 2005). Among the four AGO proteins in mammals, Argonaute RISC Catalytic Component 2 (AGO2) is the best characterized and is essential for life, as whole-body mouse knockouts of *Ago2* are embryonic lethal (Alisch et al., 2007).

Outside of its canonical cytoplasmic function in inhibition of mRNA translation (Huang and Li, 2014), several roles for AGO2 in the nucleus have been described. In *Drosophila* larvae and human fibroblasts, AGO2 mediates gene silencing by recruiting histone modifiers to target loci (Benhamed et al., 2012; Grimaud et al., 2006). AGO2 was also found to regulate alternative splicing of pre-mRNAs in *Drosophila* and HeLa cells either by coupling RNA polymerase II (RNAPII) with histone modifications such as H3K9me3, or by an alternative unknown mechanism (Ameyar-Zazoua et al., 2012; Taliaferro et al., 2013). Lastly, AGO2 was shown to facilitate DNA double strand break repair in *Arabidopsis* and cultured human cells (Wei et al., 2012). These studies point to critical nuclear roles for AGO2, but the in vivo relevance of these functions, particularly in mammals, remains poorly understood.

Spermatogenic cells offer an attractive system to reveal nuclear mechanisms of gene regulation: as mature sperm develop in mammals, the genome undergoes a unique chromatin reorganization program and employs distinct mechanisms of transcriptional regulation (Braun, 2001). The nucleus of spermatogenic cells therefore represents a valuable system where unusual mechanisms of gene regulation not evident in steady-state somatic cells may be uncovered. While *Ago2* has been studied extensively in many cellular contexts and species, its role in spermatogenesis has not been well characterized. Conditional knockout of *Ago2* in male germ cells was reported to have no phenotype, as knockout males were fertile and exhibited histologically normal spermatogenesis (Hayashi et al., 2008). However, it is unclear whether the absence of a fertility phenotype was due to low efficiency of the *TNAP-Cre* transgene used in this study, or compensation by the other AGO proteins. In contrast, conditional knockout of *Dicer* and *Drosha*, which act upstream of *Ago2* in the miRNA pathway, resulted in apoptosis, abnormal morphology, and infertility, depending on the stage of spermatogenesis that was targeted (Hayashi et al., 2008; Maatouk et al., 2008; Romero et al., 2011; Wu et al., 2012). Further, AGO4 is important for regulating entry into meiosis and silencing of sex chromosomes (Modzelewski et al., 2012), and knockout of *Ago2* in mouse oocytes led to female sterility and impaired oocyte maturation (Kaneda et al., 2009), suggesting a critical role for Argonaute proteins in gametes.

Here, we discover a role for nuclear AGO2 in positive regulation of gene expression during mammalian sperm development. This function is independent of the canonical role of cytoplasmic AGO proteins. We show that conditional knockout of *Ago2* early in spermatogenesis causes reduced sperm count and abnormal sperm head morphology, along with a decrease in protein levels of multiple genes involved in sperm head morphogenesis and motility. At the molecular level, we find that nuclear AGO2 interacts with protein partners that are distinct from its cytoplasmic interactors, including regulators of splicing, export, and chromatin remodelling. We show that AGO2 interacts with chromatin near developmentally relevant genes, binds their transcripts in the nucleus, and promotes their expression at the protein level. AGO2-chromatin interactions at these genes occur during a specific developmental interval and are temporally separated from protein-level effects, indicating that precise developmental timing is critical for this activity. These results reveal an important and unexpected role for AGO2 in positive regulation of gene expression during spermatogenic development.

## Results

### AGO2 is important for sperm development

To understand the function of *Ago2* during mammalian male reproduction, we generated *Ago2* germline conditional knockout (cKO) mice. We used a *Ddx4-Cre* transgene (Gallardo et al., 2007; Hu et al., 2013) and a conditional *Ago2* allele (O’Carroll et al., 2007) to delete exons 9-11 of the *Ago2* gene specifically in germ cells at birth. As previously reported, AGO2 protein expression was lost in spermatogenic cells following Cre activity, although the nonsense transcript was present (**Figure 1A, S1A, S1B**). We observed a significant reduction in sperm count in *Ago2* cKO males, indicating a defect in sperm production not previously reported (**Figure 1B**).

**Figure 1.**
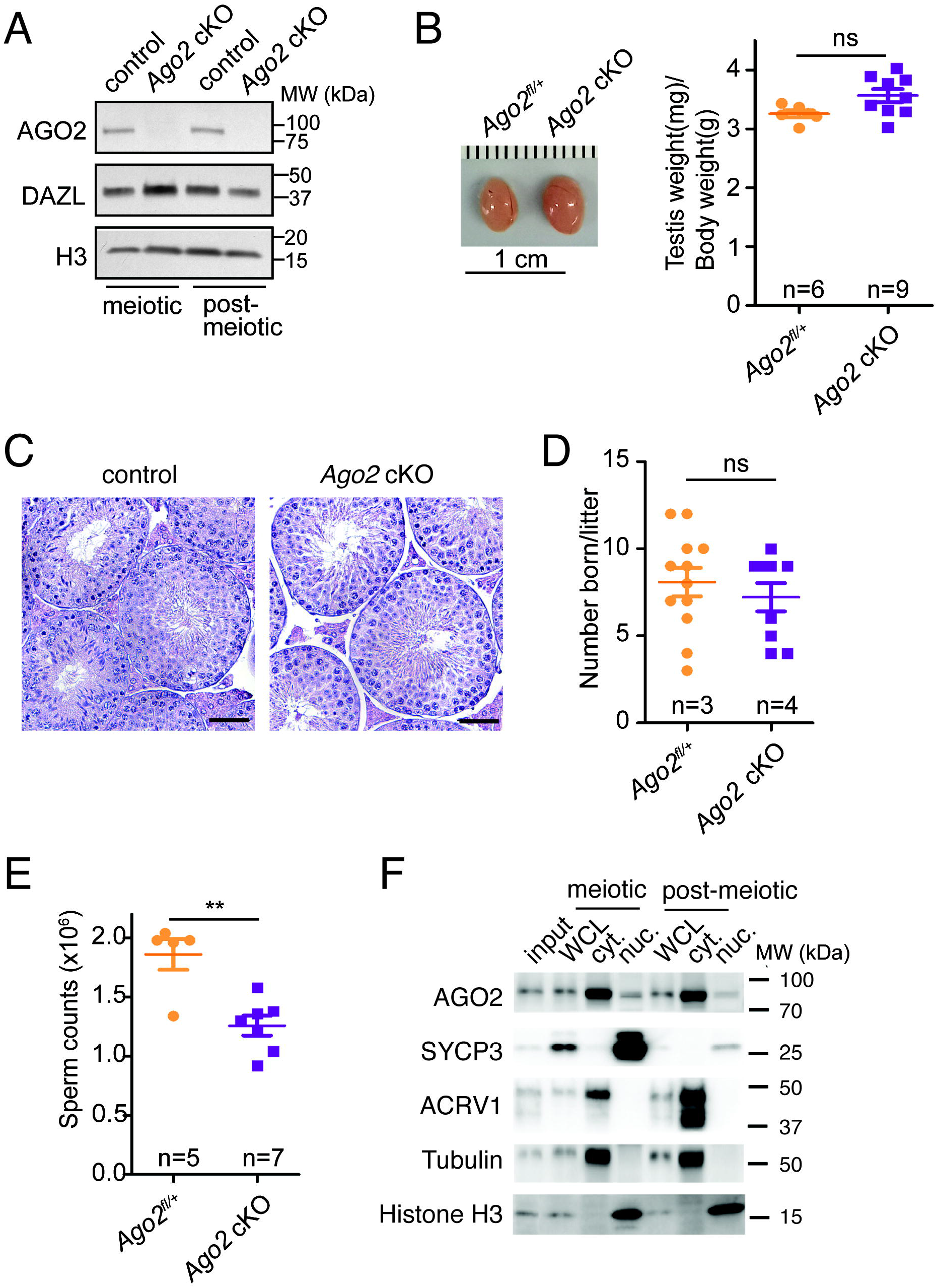
*Ago2* conditional knockout in male germ cells leads to reduced sperm count. **A**, Western blot analysis of *Ago2* knockout efficiency in meiotic and post-meiotic germ cell populations. DAZL marks germ cells; H3 is a loading control. **B**, Epididymal sperm count in *Ago2* cKO males and littermate controls. **C**, Gross images of testes from *Ago2* cKO males and littermate controls (left) and quantitation of testis/body weight ratios (right). **D**, Hematoxylin & eosin (H&E) stained testicular sections from *Ago2* cKO and control adult males. Scale bar, 50 μm. **E**, Fertility in *Ago2* cKO males and littermate controls, assessed as number of pups born per litter. **F**, Western blot analysis of AGO2 protein in whole cell lysate (WCL), cytoplasmic fractions, or nuclear fractions of meiotic (pachytene spermatocyte, SYCP3+) and post-meiotic (round spermatid, ACRV1+) male germ cells. Tubulin and histone H3 mark cytoplasmic and nuclear fractions, respectively. Error bars indicate SD. *p<0.05, **p<0.01, ***p<0.005, Welch’s t-test.

We next sought to define the developmental defects that cause reduced sperm number in the absence of AGO2. *Ago2* cKO mice exhibited no obvious change in testis size, morphology, or testis-to-body weight ratio compared to littermate controls, and spermatogenesis was grossly normal, with all spermatogenic cell types present in the seminiferous tubules (**Figure 1C, D**). There was no difference in numbers of apoptotic germ cells, and no difference in expression of transposable elements, which can cause spermatogenesis defects (Carmell et al., 2007; Pastor et al., 2014; Soper et al., 2008; Webster et al., 2005) (**Figure S1C, S1D**). *Ago2* cKO males fathered the same number of pups as littermate controls (**Figure 1E**), in agreement with the normal fertility reported by Hayashi et al in *Ago2* cKO male mice (Hayashi et al., 2008). This finding is consistent with previous reports indicating that male mice with sperm count reductions as large as 60% can still produce normal numbers of offspring under laboratory conditions (Hudson et al., 1998; Kumar et al., 1997; Santti et al., 2005; Schürmann et al., 2002; Zhang et al., 2006). Loss of AGO2 was not strongly compensated by increased transcription of other family members (**Figure S1B**). We conclude that *Ago2* is important for sperm production, although it is not required for male fertility.

As a first step toward understanding how AGO2 contributes to sperm production, we asked where AGO2 protein is expressed in male germ cells. We collected purified meiotic (pachytene spermatocyte) and post-meiotic (round spermatid) male germ cells from adult mouse testes (**Figure S1E**) and isolated nuclear and cytoplasmic fractions. We confirmed AGO2 protein expression in both nucleus and cytoplasm of meiotic and post-meiotic male germ cells, in agreement with previous observations in other mammalian tissues (Gagnon et al., 2014; Kim et al., 2011; Ravid et al., 2020) (**Figure 1F**), suggesting that AGO2 can function in both nucleus and cytoplasm at multiple steps of spermatogenic development.

### AGO2 regulates the expression of spermatogenesis genes in a stage-dependent fashion

Because AGO2 is a key protein in the miRNA and small interfering RNA (siRNA) pathways, we first postulated that AGO2 loss disrupts normal stability or translational dynamics of target mRNAs relevant to sperm maturation. To address this hypothesis, we purified meiotic and post-meiotic germ cells from *Ago2* cKO and wild type controls, and compared changes in the transcriptome and proteome of these two developmental stages in *Ago2* cKO compared to control mice using RNA-seq and quantitative mass spectrometry.

In meiotic cells, we identified 825 differentially expressed genes (DEGs) by RNA-seq, 94% of which were upregulated (**Figure 2A, Table S1**). Upregulated transcripts were enriched for several gene ontology (GO) terms related to male germ cell development and maturation (**Figure 2B**). In contrast, 321 DEGs with relatively modest effect sizes were identified in post-meiotic cells, and there was no significant bias toward up- or downregulation (**Figure 2A, Table S1**). Of these DEGs, GO analysis found an enrichment for only one term, “regulation of localization”, for downregulated transcripts, and no enriched GO terms for upregulated transcripts. There was a statistically significant overlap of 60 DEGs across both stages (30% of post-meiotic DEGs, p=1.241e-80, hypergeometric test), all of which were upregulated in both cell types. However, there was no functional enrichment for spermatogenesis or any other function in this gene set (**Figure 2C**).

**Figure 2.**
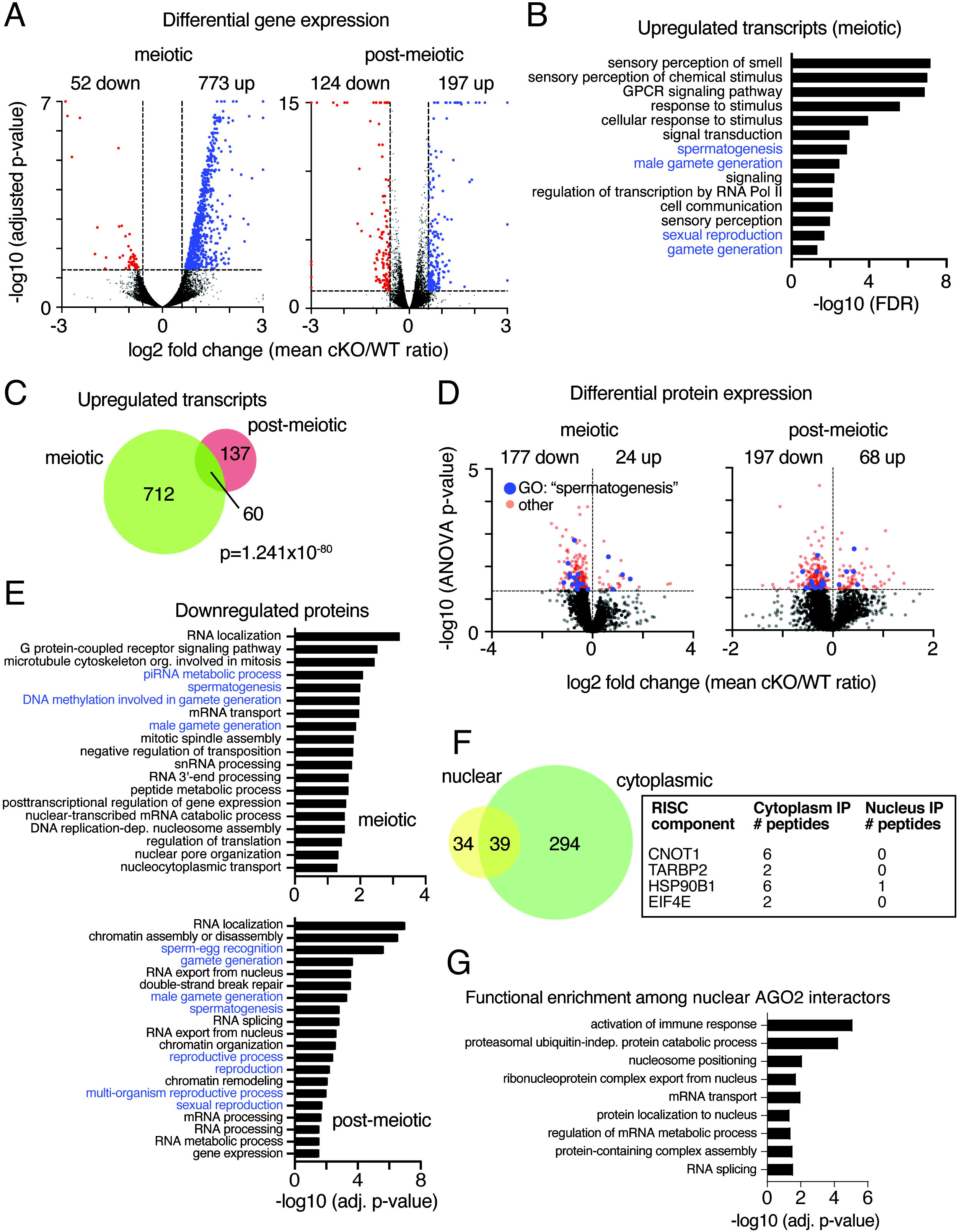
*Ago2* regulates transcript and protein expression of male reproduction-related genes in a stage-dependent fashion. **A**, Volcano plots showing global transcriptional changes in *Ago2* cKO male germ cells at meiotic (left) and post-meiotic (right) stages. Transcripts with log2 fold change ≤1.5 or ≥1.5 and adjusted p-value ≤ 0.05 are demarcated by dashed lines and highlighted in blue (upregulated) or red (downregulated). **B**, Selected enriched GO terms associated with upregulated transcripts in *Ago2* cKO meiotic cells. Terms relevant to male reproduction are highlighted in blue. **C**, Overlap of transcripts significantly upregulated in both meiotic and post-meiotic germ cells in *Ago2* cKO males. p-value, hypergeometric test. **D**, Volcano plots showing global protein level changes in meiotic and post-meiotic male germ cells from *Ago2* cKO males. Statistically significant differentially expressed proteins (p ≤ 0.05) are demarcated by a dashed line and highlighted in red. Differentially expressed proteins belonging to the GO category “spermatogenesis” (GO:0007283) are highlighted in blue. **E**, Statistically enriched GO terms associated with downregulated proteins in *Ago2* cKO meiotic and post-meiotic cells. GO terms related to male fertility are highlighted in blue. **F**, Common and compartment-specific AGO2 protein interactors in the nuclear and cytoplasmic compartments of wild type post-meiotic cells. Box shows all detected miRNA/siRNA pathway components along with the number of unique peptides identified in each compartment. **G**, Statistically enriched GO terms for AGO2 protein interactors in the nuclear compartment.

Quantification of global protein levels in *Ago2* cKO spermatogenic cells identified 201 differentially expressed proteins (DEPs) in meiotic cells and 265 DEPs in post-meiotic cells. Unexpectedly, although AGO2 usually acts as a negative regulator of protein expression, there was a strong bias towards protein downregulation in both *Ago2* cKO cell types (88% of meiotic and 74% of post-meiotic DEPs, **Figure 2D, Table S2**). In both datasets, the sets of downregulated proteins were significantly enriched for biological processes related to reproduction, along with terms related to RNA processing and chromatin organization, although only 17 specific proteins were downregulated at both stages (**Figure 2E, S2A**). Upregulated proteins were not enriched for any GO terms in meiotic cells and were weakly enriched for terms associated with the Golgi apparatus and protein processing in post-meiotic cells (**Figure S2B**). Furthermore, DEPs were poorly correlated with the DEGs identified in *Ago2* cKO cells: mRNA and protein changes were correlated for only five genes in meiotic and five genes in post-meiotic cells. Therefore, although upregulation of transcripts in *Ago2* cKO cells is consistent with the known function of AGO2 in miRNA-mediated transcript destabilization, the transcripts directly affected in germ cells do not explain the reduced sperm number and downregulation of reproduction-associated proteins observed in *Ago2* cKO males. Based on these data, the miRNA-dependent canonical function of AGO2 is unlikely to explain the abnormal sperm counts in *Ago2* cKO animals.

### AGO2 interacts with distinct protein partners in germ cell nuclei compared to cytoplasm

To further examine if the changes in mRNA and protein levels detected in *Ago2* cKO germ cells could be due to a loss of regulation by the canonical AGO2-miRNA pathway, we characterized AGO2 protein interactors by immunoprecipitation-mass spectrometry (IP-MS) in cytoplasmic and nuclear fractions of post-meiotic cells (**Figure S2C, Table S3**). As expected, four components of the miRNA/siRNA pathway (CNOT1, TARBP2, HSP90B1, and EIF4E) were detected as cytoplasmic AGO2 interactors (**Figure 2F**), along with 329 other proteins enriched for numerous biological processes including immune response and RNA localization (**Table S3**). Of the 73 proteins that interacted with AGO2 in the nucleus, more than half were also detected as cytoplasmic AGO2 interactors (**Figure 2F**). Interestingly, however, no known components of the miRNA/siRNA pathway immunoprecipitated with nuclear AGO2. Instead, the set of proteins that exclusively bound nuclear, but not cytoplasmic, AGO2 were enriched for splicing, ribonucleoprotein nuclear export, and nucleosome positioning (**Figure 2G, S2D**). The interaction of AGO2 with unique proteins in germ cell nuclei suggests that it operates via distinct pathways in the nuclear and cytoplasmic compartments.

From our transcriptome, proteome, and interactome profiles, we conclude that AGO2 is a regulator of gene expression during spermatogenesis. Our transcriptomic analysis suggests that part of this function likely involves regulation of transcript stability as part of the canonical AGO2-miRNA pathway, predominantly before or during meiosis. In contrast, protein-level effects indicate that AGO2 may also act via a separate mechanism to promote expression of a subset of spermatogenesis-related proteins. Given the presence of AGO2 in the nucleus and its distinct nuclear interactome, we hypothesized that a nuclear AGO2 pathway might be responsible for this positive regulatory function.

### AGO2 binds introns of nuclear mRNAs in meiotic and post-meiotic cells

To shed light on a possible role for nuclear AGO2 during spermatogenesis, we performed enhanced UV crosslinking and immunoprecipitation (eCLIP) to identify AGO2 binding targets in nuclei of meiotic and post-meiotic cells (Van Nostrand et al., 2016). AGO2 antibody was used to pull down RNAs from nuclear fractions. Precipitated RNAs were detected at the predicted AGO2 band (**Figure 3A**), and RNAs excised from corresponding bands were recovered to construct cDNA libraries (**Figure 3B**). 1448 RNA targets were shared between two biological replicates in meiotic nuclei, while 2006 RNA targets were shared between replicates in post-meiotic nuclei (**Table S4**). In both cell types, AGO2-bound nuclear transcripts were enriched for a variety of developmental and signaling functions (**Figure 3C, S3A**). 750 RNAs were bound by AGO2 in both meiotic and post-meiotic nuclear samples (**Table S4**), and these shared targets were enriched for a similar set of broad developmental and molecular functions (**Figure 3C, S3A**). Nuclear AGO2 may therefore have the potential to affect a wide range of physiological functions beyond spermatogenesis. Most AGO2 binding sites were located in intron regions of nuclear transcripts (62% of CLIP peaks in meiotic cells and 74% of CLIP peaks in post-meiotic cells) (**Figure 3D**). This distribution reflected the overall distribution of nucleotides in unspliced transcripts (75%) for post-meiotic cells, but was depleted in meiotic cells, where a relatively larger fraction of binding sites occurred in promoters and exons (10% vs. 6.1% for promoters and 11% vs. 4.6% for exons in meiotic CLIP targets vs. all unspliced transcripts). Most nuclear RNA targets corresponded to protein-coding genes in both meiotic and post-meiotic cells (**Figure S3B**).

**Figure 3.**
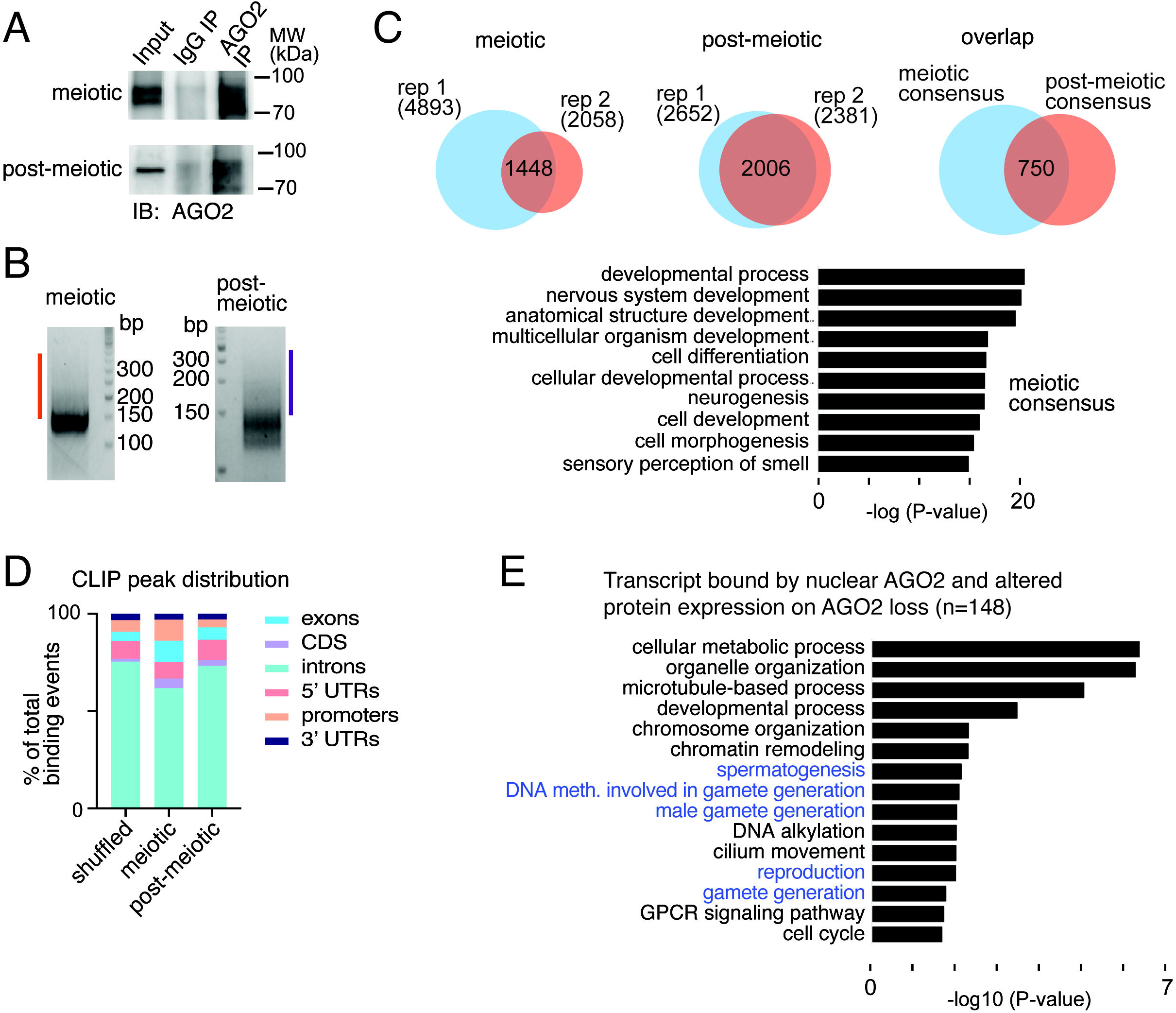
Identification of AGO2 RNA targets in nuclei of male germ cells by eCLIP. **A**, Immunoblot analysis of AGO2 immunoprecipitation. **B**, Size-selection of cDNA (orange and purple bars) following AGO2 IP, linker (147nt) ligation, and PCR amplification. **C**, Overlap of AGO2 targets from 2 eCLIP replicates from meiotic (left) and post-meiotic (center) cells, and transcripts identified in both replicates of both cell types (right). Bottom, GO analysis of meiotic consensus targets. **D**, Genomic distribution of all AGO2 eCLIP binding regions. The ‘shuffled’ control was generated by randomly shuffling a representative CLIP peak set across all annotated transcripts. **E**, Selected enriched GO terms for the sets of genes with altered protein expression and transcripts bound by AGO2 in meiotic or post-meiotic nuclei.

Of the 445 genes whose expression was altered at the protein level in either meiotic or post-meiotic germ cells (**Figure 2D**), 148 (33%) had nuclear transcripts bound by AGO2, and 122 (82%) of these were downregulated at the protein level upon loss of AGO2. Both the total set of 148 proteins and the set of 122 downregulated proteins were enriched for functions in sperm development (**Figure 3F, S3C**). Interestingly, the best correlation was observed between AGO2 mRNA binding in meiotic cells and protein expression changes in post-meiotic cells: 73 of 244 (29%) of post-meiotic differentially expressed proteins were bound by AGO2 in meiotic cells, indicating a developmental delay between nuclear transcript binding and downstream effects on cellular function. We conclude that binding of AGO2 to nuclear transcripts at meiotic stages impacts protein function, and potentially cellular phenotype, during later stages of spermatogenic development.

### AGO2 alters splicing in male germ cells and promotes nuclear export of bound transcripts

To understand the regulatory function of AGO2 on nuclear transcripts in male germ cells, we examined the sets of nuclear protein interactors identified in our IP-MS data (**Figure 2G**). We noted significant enrichment for the RNA splicing machinery, as well as several nuclear pore complex components. Nuclear AGO2 is known to associate with the splicing machinery in HeLa cells, where it promotes alternative splicing by slowing transcriptional elongation (Ameyar-Zazoua et al., 2012). We assessed differential splicing events in *Ago2* cKO compared to control in our RNA-seq datasets from both meiotic and post-meiotic germ cells using an established pipeline (Shen et al., 2012, 2014) and found that a modest number of transcripts undergo altered splicing in germ cells following loss of AGO2 (**Figure 4A, Table S5**). Splicing differences were relatively small in magnitude (**Figure 4B, S4A, S4B**), and a minority (17%) of differentially spliced transcripts were directly bound by AGO2 in our eCLIP data. Differentially spliced transcripts were not enriched for reproductive or spermatogenic functions, and of the alternatively spliced transcripts in *Ago2* cKO, there was no bias toward retention of variant exons in the absence of AGO2, contrary to the effect reported in HeLa cells in vitro (Ameyar-Zazoua et al., 2012). We conclude that, while AGO2 is involved in regulating differential splicing in male germ cells, loss of this function is unlikely to explain the reduced sperm count in *Ago2* cKO mice.

**Figure 4.**
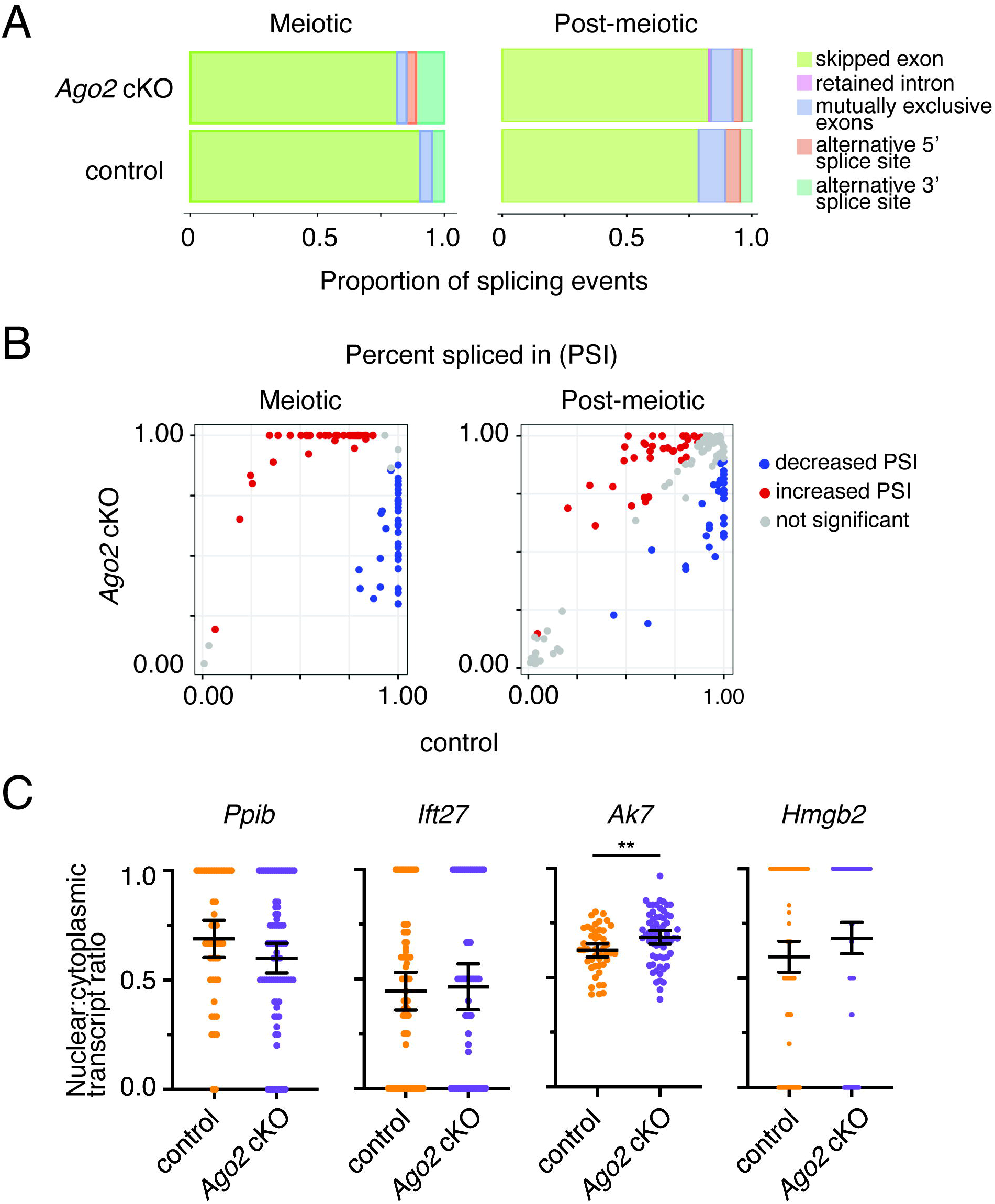
Nuclear AGO2 regulates splicing and facilitates nuclear export of mRNAs. **A**, Overall proportion of different alternative splicing events in control and *Ago2* cKO meiotic and post-meiotic germ cells. **B**, Change in exon percent spliced in (PSI) for individual transcripts with a significant difference in skipped exons in *Ago2* cKO compared to control germ cells. **C**, Nuclear:cytoplasmic ratios in post-meiotic cells for two transcripts bound (*Ak7*, *Hmgb2*) and two transcripts not bound (*Ift27*, *Ppib*) by nuclear AGO2. Localization was assessed by single-molecule RNA *in situ* hybridization (RNAscope) in *Ago2* cKO compared to control germ cells. Data points show ratios for individual cells from at least three tubules. Bars show mean and 95% confidence interval. **p<0.01, Welch’s t-test. Sample images are shown in **Figure S4C**.

Because we found enrichment of nuclear pore components among AGO2 nuclear interactors, as well as the nuclear export factor NXF1 (**Fig. S2D**), we next asked whether AGO2 might facilitate export of transcripts from nucleus to cytoplasm. We selected two transcripts bound by nuclear AGO2 that were also depleted at the protein level in *Ago2* cKO germ cells (*Ak7* and *Hmgb2*), along with two negative control transcripts not bound by nuclear AGO2 (*Ift27* and *Ppib*), and evaluated nuclear/cytoplasmic transcript distribution semiquantitatively using RNAscope (**Figure 4C, S4C**). We found a modestly increased ratio of nuclear to cytoplasmic transcripts for the AGO2-bound transcript *Ak7*, indicating increased nuclear retention. *Hmgb2* transcripts also showed a trend toward increased nuclear retention, although the difference was not statistically significant (p=0.0949, Welch’s t-test). Neither control gene had increased fractions of nuclear transcripts in *Ago2* cKO meiotic cells (**Figure 4C, S4C**). We conclude that AGO2 binding of nuclear transcripts promotes export of associated mRNAs, possibly through interaction with the nuclear pore and NXF1, although this effect is relatively modest.

### AGO2 interacts with open chromatin in meiotic male germ cells

Because AGO2 has previously been shown to associate with chromatin and directly regulate transcription (Cernilogar et al., 2011; Moshkovich et al., 2011; Nazer et al., 2018), we next evaluated AGO2-chromatin interactions in spermatogenic cells. We performed chromatin immunoprecipitation followed by sequencing (ChIP-seq) for AGO2 in pre-meiotic (differentiating spermatogonia), meiotic, and post-meiotic cells, after validating our ChIP protocol in mouse embryonic stem cells (mESCs) (**Figure S5A, Table S6**). In mESCs and in pre- and post-meiotic germ cells, we found relatively few, but strong, peaks (148 peaks in mESCs, 164 peaks in pre-meiotic cells, and 142 peaks in post-meiotic cells). In contrast, we identified thousands of broader, weaker peaks in meiotic cells (**Figure 5A, 5B, Table S7**). Notably, peaks were frequently shared across pre- and post-meiotic germ cells, but these locations rarely overlapped with peaks found in meiotic cells (**Figure 5A, 5B, S5B, Table S7**). Meiotic peaks were more frequently intergenic, whereas pre-meiotic and post-meiotic peaks were most often found in gene bodies (**Figure 5C**). AGO2 ChIP-seq in *Ago2* cKO germ cells yielded significantly reduced or absent signal compared to control, confirming that the detected binding sites were specific to AGO2 (**Figure 5B, S5B, S5C**). Together, these data suggest that AGO2 interacts with chromatin in a developmentally dynamic manner during spermatogenesis.

**Figure 5.**
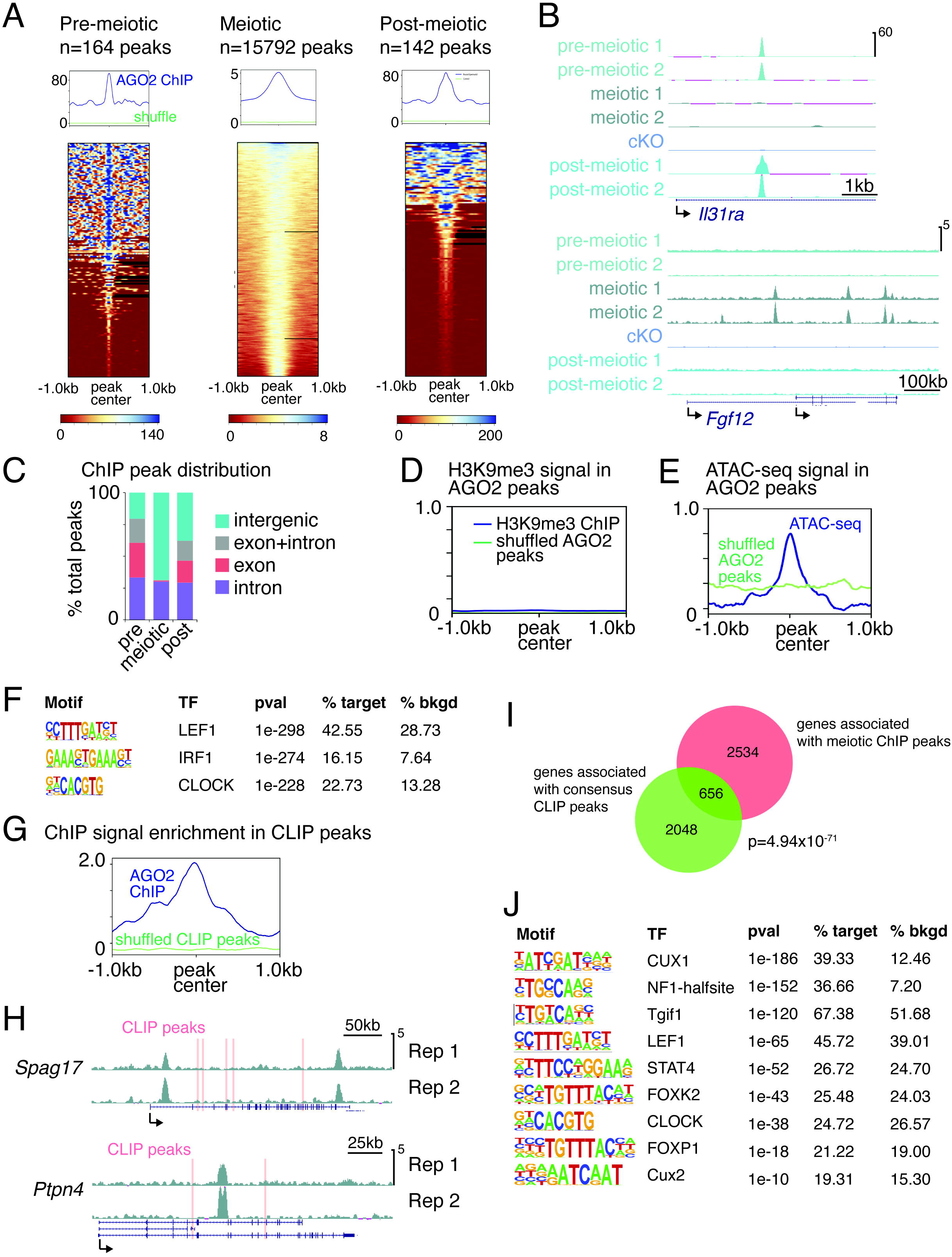
AGO2 interacts with chromatin at genes with AGO2-bound nuclear transcripts in a developmentally dynamic manner. **A**, Enrichment of AGO2 on chromatin in pre-meiotic, meiotic, and post-meiotic male germ cells. Top, metagenes showing average ChIP signal at AGO2 peaks, with reference point set to the peak center. ‘Shuffle’ shows ChIP signal when the same number of size-matched peaks were randomized across the genome. Bottom, heatmaps showing ChIP signal enrichment at each peak. **B**, Genome browser tracks showing representative AGO2 ChIP peaks in two replicates from each cell type, and from *Ago2* cKO mixed meiotic and post-meiotic cells. **C**, Distribution of AGO2 peaks in introns, exons, and intergenic regions. **D**, Metagene of H3K9me3 ChIP signal at AGO2 peaks in meiotic cells. **E**, Metagene of ATAC-seq signal at AGO2 peaks in meiotic cells. **F**, Top motifs enriched in AGO2 ChIP peaks in meiotic cells. **G**, Ago2 ChIP enrichment signal at eCLIP peaks in meiotic cells relative to shuffled eCLIP peaks. **H**, Genome browser tracks showing representative genes associated with both ChIP and eCLIP peaks. **I**, Overlap between genes associated with ChIP peaks in meiotic cells and genes containing eCLIP peaks in both meiotic and post-meiotic cells. p-value, hypergeometric test. **J**, Motifs enriched in AGO2 ChIP peaks associated with genes that also contain eCLIP peaks. Only motifs matching a transcription factor expressed in meiotic cells are shown.

We next sought to determine if AGO2-chromatin interactions correlate with the transcriptional state of the nearby genome. AGO2 and other AGO family proteins can associate with H3K9me3 and heterochromatin in other organisms (Ameyar-Zazoua et al., 2012; Verdel et al., 2004; Zilberman et al., 2003). However, H3K9me3 was not associated with AGO2 binding sites in our data (Liu et al., 2019) (**Figure 5D**), and we found no change in global levels of H3K9me3 in *Ago2* cKO meiotic or post-meiotic cells (**Figure S5D**). Given the lack of enrichment in heterochromatin, we then tested whether AGO2 binding sites were preferentially found in regions of open chromatin. We evaluated ATAC-seq signal strength at AGO2 peaks (Maezawa et al., 2018) and found a moderate enrichment for open chromatin (**Figure 5E**). In meiotic cells, open chromatin is found at regions of transcriptional activity, but also at sites of recombination between homologous chromosomes (Baker et al., 2014; Lange et al., 2016). To distinguish between these types of open chromatin, we then asked if AGO2 in meiotic cells preferentially localized to recombination hotspots. We compared our AGO2 data to a published set of mouse recombination hotspots (Smagulova et al., 2016) and found that only 15 meiotic AGO2 peaks overlapped hotspots, compared to 280 overlaps when AGO2 peaks were randomized across the genome. This finding indicates no significant association of AGO2 with recombination hotspots and suggests that AGO2 preferentially binds to the genome near sites of open chromatin corresponding to transcriptional activity in meiotic cells.

Because we found that AGO2 binds nuclear mRNAs (**Figure 3**) and associates with open chromatin, we wondered if its association with chromatin was dependent on interaction with nascent transcripts. We addressed this question using mESCs, due to the limited cell numbers available for spermatogenic cells. We treated mESC nuclei with an RNAse cocktail to cleave any nascent transcripts, and evaluated AGO2 interaction intensity in RNAse-treated cells (**Figure S5E**). AGO2 binding did not change significantly in RNAse-treated cells relative to control, suggesting that AGO2 interaction with chromatin does not require RNA transcripts.

To understand how AGO2 is recruited to meiotic chromatin, we defined motifs enriched in AGO2 ChIP peaks in meiotic cells. Several enriched motifs matched binding sites for transcription factors expressed during male meiosis (**Figure 5F, S5F**), suggesting that one or more of these transcription factors may recruit AGO2 to target sites, similar to transcription factor-mediated recruitment of AGO2 to chromatin in other systems (Benhamed et al., 2012; Cernilogar et al., 2011; Moshkovich et al., 2011; Nazer et al., 2018). However, none of the transcription factors matching enriched motifs was identified as a nuclear AGO2 interactor in our IP-MS dataset, implying that recruitment may be indirect or mediated by factors not identified in our analysis.

### AGO2 binds chromatin near hundreds of genes corresponding to AGO2-bound nuclear transcripts in meiotic cells

To determine the functional significance of AGO2-chromatin interactions in spermatogenic cells, we next compared our ChIP and eCLIP datasets from meiotic cells to ask if there was a relationship between genomic sites and nuclear transcripts bound by AGO2. Analysis of AGO2 ChIP-seq signal intensity at CLIP peaks relative to shuffled control demonstrated a moderate enrichment for AGO2 binding at genomic sites corresponding to AGO2-bound mRNA transcripts in meiotic but not post-meiotic cells (**Figure 5G, S5G**). The relatively weak level of enrichment seen in CLIP peak locations could be explained by variability in the exact placement of the ChIP peak relative to the CLIP peak when both occur near the same gene. Indeed, we consistently observed maximum ChIP signal near, but not directly overlapping, a CLIP peak (**Figure 5H, S5H**). After overlapping the sets of genes associated with ChIP and CLIP peaks, we found a significant overlap of 656 genes for which AGO2 binds chromatin at or near (<20kb) the coding region and also binds the nuclear transcript (**Figure 5I, Table S8**). ChIP peaks associated with these genes were enriched for nine motifs matching transcription factors expressed in male meiotic germ cells (**Figure 5J**), further refining a set of candidate protein partners that may mediate AGO2 function on chromatin. We predicted that changes in expression of proteins encoded by this set of 656 target genes could mediate the reduction in sperm count observed in *Ago2* cKO males.

### Loss of *Ago2* causes sperm head defects and impaired expression of proteins required for sperm morphogenesis

Multiple genes required for sperm morphology and motility were bound by AGO2 at the chromatin and transcript levels in the nucleus and downregulated at the protein level in the *Ago2* cKO, and decreased sperm count is often associated with sperm morphology and sperm motility defects (Macleod, 1971; Zhuang et al., 2014). We therefore evaluated motility and morphology in *Ago2* cKO sperm. Using computer assisted semen analysis (CASA), we found a 7% increase in the fraction of slow-moving sperm among *Ago2* cKO cauda epididymal sperm after capacitation (functional sperm maturation), but no significant difference in overall progressive motility compared to control (**Figure S6A, S6B**); we conclude that loss of AGO2 has at most a minor effect on motility. We then tested for sperm morphology defects using Coomassie staining of cauda epididymal sperm and blinded scoring of sperm head shape in *Ago2* cKO and control animals. *Ago2* cKO sperm had a significantly higher proportion of head morphology defects compared to control (∼37% vs. ∼3% abnormal), predominantly in the form of small, rounded heads (**Figure 6A**). Tail lengths were comparable between *Ago2* cKO and control sperm (**Figure S6C**). Defects in formation of the acrosome, the organelle containing enzymes that break down the zona pellucida at fertilization, are the major cause of sperm head defects (Perrin et al., 2013), and evaluation of acrosome formation by lectin-peanut agglutinin staining showed that the acrosome was small and improperly formed in *Ago2* cKO sperm compared to controls (**Figure 6B**). The set of head morphology defects observed in *Ago2* cKO sperm parallels the defects expected following depletion of several of the proteins downregulated in *Ago2* cKO post-meiotic germ cells, including HMGB2, YBX2, and PTBP2 (Catena et al., 2006; Hannigan et al., 2018; Lorès et al., 2018; Snyder et al., 2015). Further, at least two of the transcription factors with binding motifs enriched among the set of dual AGO2 ChIP and CLIP targets (NF1 and STAT4; **Figure 5J**) are known to regulate sperm head formation (Chohan et al., 2018; Herrada and Wolgemuth, 1997). Phenotype enrichment analysis (Weng and Liao, 2017) confirmed that downregulated DEPs were statistically significantly (adjusted P-value ≤0.05) enriched for abnormal male germ cell morphology and other reproductive phenotypes including male infertility and arrest of spermatogenesis. We further validated downregulation of five of these genes by Western blotting in *Ago2* cKO post-meiotic cells (**Figure 6C).**

**Figure 6.**
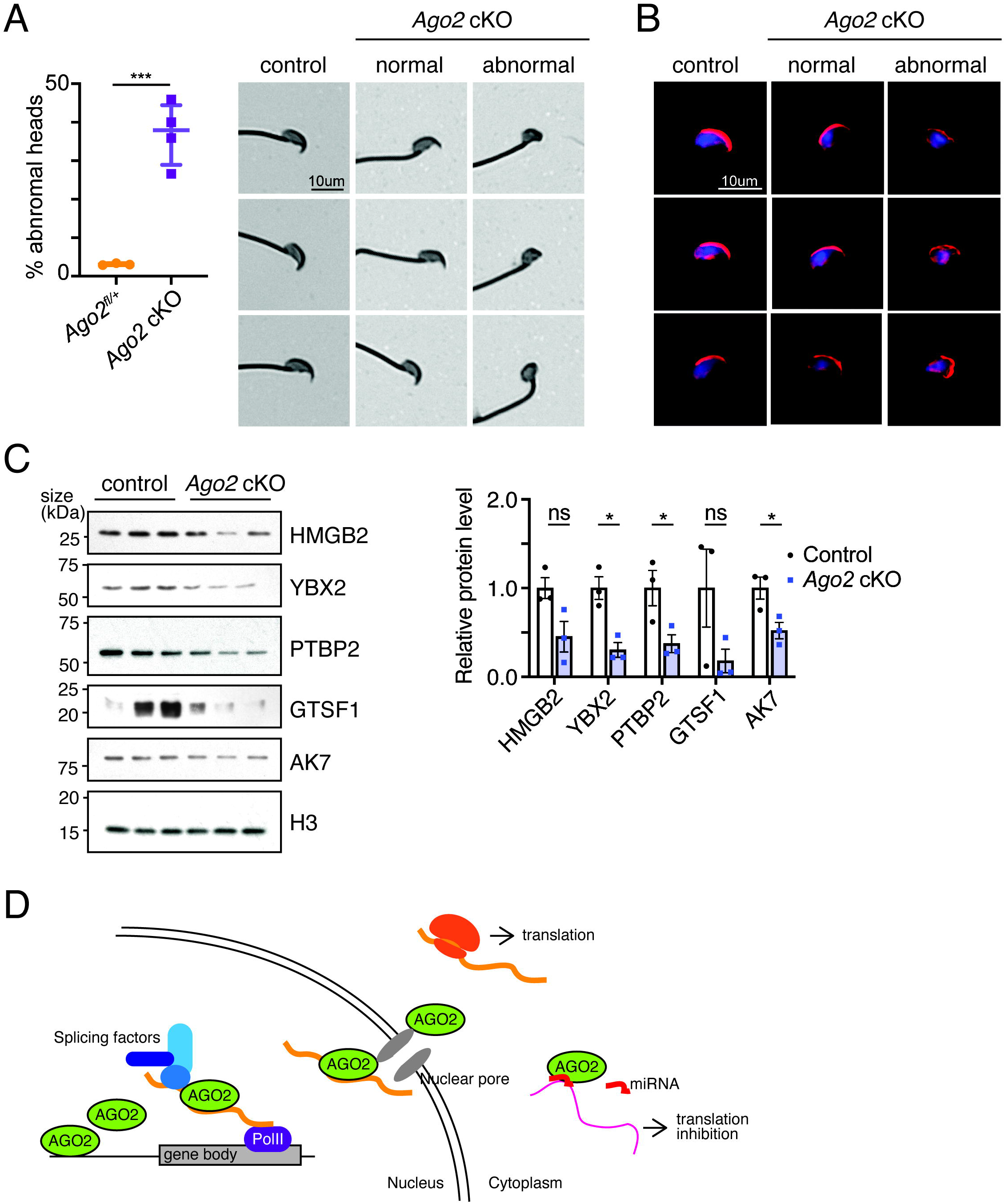
Loss of AGO2 impairs expression of proteins required for normal sperm production and results in morphologically abnormal sperm. **A**, Left, percentage of *Ago2* cKO spermatozoa with abnormal heads, representing n>400 spermatozoa from each of 3-4 biological replicates. Mean of each replicate is shown along with the median and interquartile range of all replicates. ***p < 0.001, Welch’s t-test. Right, brightfield images of spermatozoa stained with Coomassie Blue, showing representative cells collected from control and *Ago2* cKO cauda epididymides. Both normal and abnormal examples of *Ago2* cKO sperm are shown. **B**, Representative spermatozoa stained with fluorophore-conjugated lectin-peanut agglutinin to visualize the acrosome (red) and DAPI to show DNA (blue). Both normal and abnormal examples of *Ago2* cKO sperm are shown. **C**, Western blot validation of downregulation of proteins involved in spermiogenesis. Bar chart shows levels of each protein relative to histone H3, quantified using densitometric analysis for three biological replicates per group. Error bars = SEM, ns, not statistically significant, *p ≤0.05, Welch’s t-test. **D**, Model for RISC-independent nuclear AGO2 function in germ cells. AGO2 interacts with chromatin to facilitate binding to nearby nascent transcripts, where it interacts with the splicing machinery. AGO2 then maintains its interaction with the spliced transcript and helps to shuttle these transcripts to the cytoplasm through its interaction with components of the nuclear pore and export machinery. Once in the cytoplasm, a subset of these transcripts may also be regulated by canonical miRNA-dependent AGO2 machinery.

We conclude that depletion of AGO2 nuclear targets at the protein level can explain the sperm phenotypes observed in *Ago2* cKO males, and that AGO2 acts on nuclear mRNAs after transcription to enforce expression of proteins required for spermiogenesis. Loss of *Ago2* in the male germline therefore undermines sperm production despite the fertility of *Ago2* cKO males under laboratory conditions, and may impair the ability of sperm to withstand genetic and environmental perturbations.

## Discussion

We examined the role of AGO2 in male mammalian germ cells. We found that AGO2 binds chromatin and nuclear transcripts in a developmentally dynamic manner during spermatogenesis. Nuclear AGO2 does not act through an association with the RISC complex, in contrast to its cytoplasmic function (Gurtan and Sharp, 2013), but instead interacts with a variety of other protein partners, including factors involved in splicing, mRNA export, and nucleosome positioning. AGO2 binds open regions of the genome and associates with both DNA and nuclear mRNA for hundreds of genes in vivo. Several of these genes are required for spermiogenesis, specifically sperm head development and acrosome formation, and loss of AGO2 results in formation of morphologically abnormal sperm and decreased total sperm count, implicating nuclear AGO2 function in regulation of these processes. Notably, the decrease in protein levels of spermatogenesis-associated genes following *Ago2* knockout is opposite to the expected effect based on the canonical role of AGO2 in miRNA-mediated translation inhibition.

Based on our findings, we propose a model (**Figure 6D**) wherein nuclear AGO2 positively regulates expression of developmentally important genes in the mammalian male germline, contrasting with the canonical function of cytoplasmic AGO2. AGO2 undergoes widespread, transient recruitment to euchromatin during prophase of meiosis, where it binds nearby nascent transcripts and promotes their export from the nucleus. AGO2-mediated mRNA regulation may therefore begin prior to its canonical miRNA-mediated role in the cytoplasm.

Our data cannot completely rule out an indirect effect of cytoplasmic AGO2 on positive regulation of protein expression. miRNA-mediated downregulation of translation-enhancing factors could cause secondary downregulation of target proteins in post-meiotic cells. However, we did not detect downregulation of translation factors at the protein level in our data. Our AGO2 IP-MS data revealed an interaction between cytoplasmic AGO2 and several translation initiation factors, raising the alternative possibility that depletion of AGO2 could destabilize this complex and reduce overall translation efficiency, although levels of these factors were also unchanged at the protein level in our quantitative proteomics dataset. Nevertheless, future experiments where AGO2 is excluded from the nucleus, but unaltered in the cytoplasm, will be required to definitively rule out cytoplasmic AGO2 function as an explanation for the sperm phenotype observed in the *Ago2* cKO.

Additional work will also be needed to fully define the molecular mechanism by which AGO2 promotes protein expression for its nuclear targets. Loss of AGO2 did not have a major effect on transcriptional elongation or splicing in male germ cells, in contrast to previous reports implicating nuclear AGO2 in these processes (Ameyar-Zazoua et al., 2012; Nazer et al., 2018; Tarallo et al., 2017). We found that AGO2 in germ cells interacts with the nuclear export machinery, and loss of AGO2 does affect the relative fraction of nuclear compared to cytoplasmic transcripts for spermiogenesis-associated genes. Therefore, nuclear AGO2 may facilitate export of target mRNAs, regulating protein abundance promoting access to translation machinery in the cytoplasm. In addition, we found several strongly enriched motifs in AGO2 ChIP peaks, implying that recruitment of AGO2 to the genome may be mediated by protein-protein interactions with transcription factors or transcription factor complexes, and these transcription factors may also modulate AGO2 function on chromatin and interaction with target transcripts.

Canonically, cytoplasmic AGO2 associates with miRNAs to buffer gene expression against perturbations to normal cell function (Ebert and Sharp, 2012; Hornstein and Shomron, 2006; Strovas et al., 2014). The nuclear function of AGO2 function we identify here may also contribute to maintaining germline function in the context of environmental stressors, such as temperature or chemical exposure, and internal stressors, such as aging or inflammatory activity. If so, we predict that the relatively mild spermatogenesis phenotype found in *Ago2* cKO mice would be enhanced under stress conditions. Alternatively, the mild phenotype could be explained by partial compensation of AGO2 loss by other members of the AGO family. Simultaneous depletion of multiple AGO family members in the male germline will help to answer this important question.

Our data indicate that nuclear AGO2 activity in the male germ line occurs primarily during meiotic prophase, a cellular state unique to germ cells. Is this nuclear function specific to the male germ line, or does it also occur during developmental transitions in other cell types? On one hand, gene regulation in germ cells often involves unique mechanisms. We observed a delay between meiotic AGO2 nuclear activity and post-meiotic effects on protein expression and phenotype, a common phenomenon in male germ cells, where transcription shuts down during post-meiotic sperm maturation and post-transcriptional regulation is required to temporally control protein expression during this stage (Rathke et al., 2014; Steger et al., 1998). On the other hand, chromatin reorganization during meiosis may reveal general regulatory functions that are relevant but not immediately apparent in other cell types. Supporting a more generalizable role for nuclear AGO2 is the fact that many of its gene and transcript targets are involved in somatic, rather than spermatogenic, development. In addition, sixteen of the nuclear AGO2 protein partners we identify here were recently found to interact with AGO2 in nuclei of mouse neurons (Ravid et al., 2020). AGO2 may therefore regulate a common set of genes across multiple cell types, with tissue-specific functions imposed by additional downstream regulatory mechanisms. It will be important to further elucidate the biological and molecular significance of nuclear AGO2 function both in the male mammalian germ line and in somatic cells.

## Acknowledgments

We thank P. Reddi for gift of the ACRV1 antibody, J. Hwang for help with STA-PUT, Y. Kong for assistance with eCLIP analysis, and S. Nicoli for critical reading of the manuscript. We thank Florine Collin from the MS & Proteomics Resource at Yale University for assistance with the proteomics sample preparation and data collection. We greatly appreciate help from the Yale Center for Genome Analysis for high-throughput sequencing. Proteomics data were collected on a mass spectrometer supported by NIH SIG S10OD018034 and Yale School of Medicine. We acknowledge start-up funds from Yale University School of Medicine and funding from the National Institute of Child Health and Human Development to J.-J.C. (R01HD096745) and to B.J.L (R01HD098128), and support from the Searle Scholars Program to B.J.L. Bluma J. Lesch, M.D., Ph.D., holds a Career Award for Medical Scientists from the Burroughs Wellcome Fund.

## Author Contributions

Conceptualization, K.N.G., H.L., B.W.W., and B.J.L.; Validation, B.W.W.; Formal Analysis, K.N.G., H.L., B.W.W., T.L. and B.J.L.; Investigation, K.N.G., H.L., B.W.W., H.W., C.B.K., J.K., and T.L.; Resources, T.L., J.-J.C., and B.J.L.; Writing – Original Draft, K.N.G., H.L., and B.W.W.; Writing – Review & Editing, A.L.C., J.-J.C., and B.J.L.; Visualization, K.N.G., H.L., and B.W.W.; Supervision, B.J.L., Funding Acquisition, J.-J.C. and B.J.L.

## Methods

### Antibodies

ChIP-seq and eCLIP-seq were performed using a mouse monoclonal antibody to AGO2 (Millipore, 04-642). This antibody has been validated for ChIP-seq applications in human (Woolnough et al., 2015). Immunoblotting was performed using the following primary antibodies and dilutions: Histone H3 (Abcam, ab1791, 1:40000 or Abcam, ab18521, 1:1000), AGO2 (Abcam, ab186733, 1:2000 or Abcam, ab32381, 1:1000), H3K9me3 (Abcam, ab8898, 1:4000), HMGB2 (Abcam, ab124670, 1:3000), DAZL (Abcam, ab34139, 1: 2000), YBX2 (Abcam, ab154829, 1:4000), AK7 (Novus Biologicals, NBP2-92176, 1: 3000), GTSF1 (NOVUS Biologicals, NBP1-83934,1:1000), PTBP2 (Proteintech, 55186-1-AP, 1:4000), α-TUBULIN (Santa Cruz, sc-8035, 1:1000), SYCP3 (Abcam, ab97672, 1:1000), ACRV1 (guinea pig, gift of Dr. Prabhakara Reddi, University of Illinois (Osuru et al., 2014), 1:5000).

### Primers

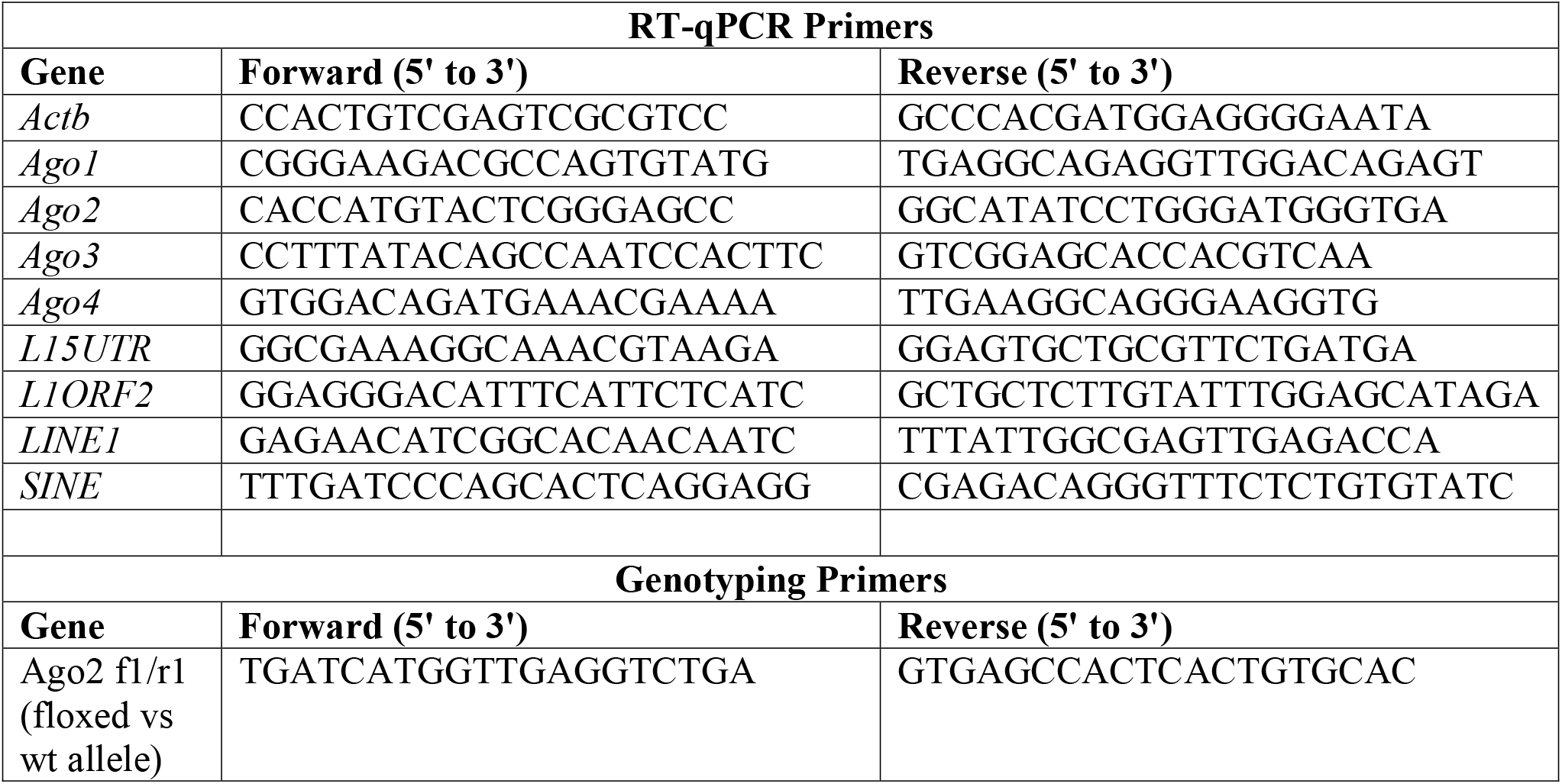
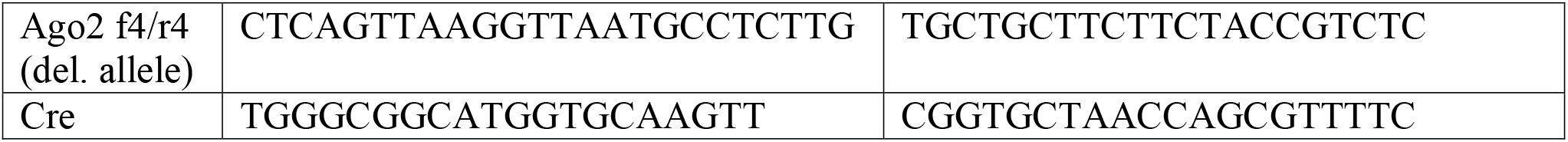

### Cell Culture

Mouse embryonic stem cells were grown at 37°C and 5% CO_2_ on 0.1% gelatin and feeder cells (Gibco CF-1 MEF MITC-Treated) in DMEM (Gibco 11965-092) + 15% fetal bovine serum (Sigma-aldrich F2442), 1:100 pen-strep (Gibco 15140-122), 1:100 GlutaMax (Gibco 35050-061), 1:100 MEM NEAA (Gibco 11140-050), Sodium Pyruvate 1:100 (Gibco 11360-070), HEPES (Gibco 15630-080), β-mercaptoethanol (Sigma-Aldrich M6250), and 1:10000 LIF (Millipore ESG1106). Cells were passaged 1:10 every 2–3 days with 1× trypsin-EDTA 0.25% (Gibco 25200-056).

### Mouse Strains and Animal Care

All mice were maintained on a C57BL/6 genetic background. *Ago2*^flox/flox^ mice (*B6.129P2(129S4)-Ago2^tm1.1Tara^/J*, (O’Carroll et al., 2007)) were obtained from Jackson Laboratory (strain #016520). *Ago2*^flox/flox^ mice were mated with mice carrying one copy of the *Ddx4-Cre* transgene (B6-*Ddx4^tm1.1(cre/mOrange)Dcp^*, (Gallardo et al., 2007; Hu et al., 2013)) in which Cre is expressed specifically in germ cells beginning at embryonic day 15.5, to generate *Ago2* conditional knockout males (*Ddx4*-*Cre*/+*;Ago2*^flox/⊗^). Adult wild type mice were purchased from Taconic Biosciences. All mice used in these studies were maintained and euthanized according to the principles and procedures described in the National Institutes of Health Guide for the Care and Use of Laboratory Animals. These studies were approved by the Yale University Institutional Animal Care and Use Committee under protocol 2020-20169 and conducted in accordance with the specific guidelines and standards of the Society for the Study of Reproduction.

### Testis collection and cell dissociation

Seminiferous tubules were isolated by enzymatic digestion (0.75-1 mg/ml collagenase IV, Gibco 17104-019) for 10 min in a 37°C water bath as described previously (Bryant et al., 2013). Male germ cells were obtained from seminiferous tubules using a second enzymatic digestion (0.25% trypsin (EDTA-free, Sigma), 2 mg/ml DNase I (Stem Cell Technologies, 07900)). The trypsin was neutralized with DMEM containing 10% fetal bovine serum (Life Technologies) and removed by centrifugation at 400xg for 5 mins before resuspending. The cell suspension was filtered in two steps through a 100 μm followed by a 40 μm cell strainer, and then used for either STA-PUT or flow cytometry.

### STA-PUT

STA-PUT was performed as described previously (Bellve, 1993). For each isolation, 6-8 testes from adult wild type mice were pooled and washed twice in Dulbecco Modified Eagle medium (DMEM). A linear gradient was generated using 350 ml of 2% BSA and 350 ml of 4% BSA solutions in the corresponding chambers. Approximately 1 × 10^8^ male germ cells were resuspended in 20ml of 0.5% BSA solution and loaded to the sedimentation chamber. After 3 hours of sedimentation in the sedimentation chamber, 60 fractions were collected in 15 ml centrifuge tubes and numbered sequentially 1-60. Cells from each fraction were collected by centrifugation at 500xg for 5 min and resuspended in 0.2 ml total volume. An aliquot of each fraction was stained with Hoechst dye (Invitrogen, H3570) and examined thoroughly by eye under phase-contrast and fluorescence microscopes to assess cellular integrity and identify cell types. Fractions containing >80% cells of appropriate size and morphology were pooled as pachytene spermatocytes (‘meiotic cells’) and round spermatids (‘post-meiotic cells’).

### Flow cytometry

For flow sorting, dissociated cells were diluted to 5 x 10^6^ cells/ml. For isolation of round spermatids, the cell suspension was incubated at 37°C with 2 µl/ml Vybrant DyeCycle Green (Life Technologies, V35004) for 20 minutes. Round spermatids were sorted based on 1C DNA content and size using a 2-laser sorter (BioRad S3e with 488nm and 561nm lasers) gated on single cells as previously described (Lesch et al., 2019). Differentiating spermatogonia (‘pre-meiotic cells’) were isolated as c-Kit+ spermatogenic cells using anti-c-Kit antibody directly conjugated to PE (eBioscience <u>12-1171-82</u>).

### Morphological assessment of spermatozoa

Cauda epididymides were dissected from adult (8-16 week old) male mice and allowed to swim out into PBS in a 12 well culture dish for 15 mins. The sample was then centrifuged for 2 min at 2000xg at 4°C and the resulting pellet was resuspended in 4% paraformaldehyde for 20 mins at room temperature. Fixed spermatozoa were pelleted and washed twice in 100 mM ammonium acetate, then resuspended in a final volume of 100 µl wash buffer. 7 µl of this cell suspension was smeared onto a glass slide and air dried followed by staining for 10 min in Coomassie blue solution or for 30 mins in 10ug/ml fluorescent dye-conjugated lectin-peanut agglutinin (lectin PNA, Invitrogen, L21409) in PBS. Coomassie stained slides were rinsed in water and allowed to air dry before coverslipping with Permount (Fisher Scientific, SP15-100). The proportion of sperm with abnormal heads was visually assessed from >400 sperm for each of 3-4 biological replicates using brightfield microscopy under blinded conditions. Slides stained with lectin PNA were rinsed in PBS, coverslipped with ProLong Gold Antifade Mountant with DAPI (Invitrogen, P36931) and imaged using confocal microscopy. Statistical significance was tested by an unpaired t-test with Welch’s correction.

### Sperm motility assessment

Cauda epididymal sperm were collected by swim-out in HEPES buffered saline (HS) medium. Aliquots of sperm were placed in a slide chamber (CellVision, 20 mm depth) and motility was examined on a 37°C stage of a Nikon E200 microscope under a 10X phase contrast objective (CFI Plan Achro 10X/0.25 Ph1 BM, Nikon). Images were recorded (40 frames at 50 fps) using a CMOS video camera (Basler acA1300-200um, Basler AG, Ahrensburg, Germany) and analyzed by computer-assisted sperm analysis (CASA, Sperm Class Analyzer version 6.3, Microptic, Barcelona, Spain). Sperm total motility and hyperactivated motility was quantified simultaneously. Over 200 motile sperm were analyzed for each trial. Three and two biological replicates were performed for control and *Ago2* cKO, respectively.

### Nuclear/cytoplasmic fractionation

A pellet of 0.5-2×10^7^ fresh spermatogenic cells was resuspended in 0.5-1 ml Cell Lysis Buffer 1 (50 mM Tris-HCl pH 7.4, 100 mM NaCl, 0.1% NP-40, with freshly added Protease Inhibitor Cocktail) and incubated on ice for 20 min with inversion every 4 min. Cell pellets were then spun down at 1200xg for 5 min at 4°C, and supernatants were kept as cytoplasmic lysate. Pellets were washed once with 1ml of Cell Lysis Buffer 2 (50 mM Tris-HCl pH 7.4, 100 mM NaCl, 0.2% NP-40) and once 1ml of Cell Lysis Buffer 3 (50 mM Tris-HCl pH 7.4, 100 mM NaCl, 0.5% NP-40), and incubated on ice for 5 min each time. Finally, the nuclear pellets were spun down at 1200xg for 5 min at 4°C, supernatants were discarded and pellets were resuspended with 500 μl of 1 ml of iCLIP lysis buffer (50 mM Tris-HCl pH 7.4, 100 mM NaCl, 1% NP-40, 0.1% SDS, 0.5% sodium deoxycholate) and incubated on ice for 5 min. Lysates were centrifuged at 15000xg for 15 min at 4°C, and the supernatants were collected as nuclear lysate.

### Immunoblot analysis

2 μg of total protein derived from meiotic or post-meiotic germ cell populations was loaded onto a Mini-PROTEAN TGX gel (Bio-Rad, 456-8093) and separated by SDS-PAGE for 1 hour at 150V. Separated proteins were then wet transferred to a 0.45 μm nitrocellulose membrane (GE Healthcare Life Science, 10600003) in Towbin buffer (25mM Tris, 192mM glycine, 20% (v/v) methanol) at a constant current of 250mA for 1 hour. Following transfer, membranes were briefly incubated in Ponceau S (Sigma Aldrich, P7170) to assess transfer quality. Membranes were then blocked in 5% w/v non-fat dry milk-TBST for 30 min with gentle agitation at room temperature. Primary antibodies were diluted in blocking buffer and incubated with the membrane overnight at 4°C with gentle rocking. Three 3-minute washes were performed with TBST before incubating immunoblotted membranes for 1 hour at RT with peroxidase-conjugated anti-rabbit secondary antibody (Jackson Immuno Research, 111-035-003) diluted 1:20 000 in TBST. Following washing, the membrane was incubated in Pierce ECL substrate (Thermo Fisher, 32109) for 5 min at RT and the signal was captured with X-ray film (Thermo Fisher, 34090) for 5-60 sec. Immunoblots were quantified by densitometry using FIJI software (Schindelin et al., 2012). Unpaired t-tests were performed for each protein to assess statistical significance.

For Western blots of CLIP samples, protein samples were prepared by iCLIP lysis buffer (50 mM Tris-HCl pH 7.4, 100 mM NaCl, 1% NP-40, 0.1% SDS, 0.5% sodium deoxycholate) containing a protease inhibitor cocktail (Roche, 11697498001). Protein samples were separated by SDS-PAGE and blotted onto the PVDF membranes. Transferred membranes were blocked in 5% w/v non-fat dry milk for 1h at room temperature, followed by incubation with diluted primary antibodies at 4°C overnight. After three washes with 0.05% TBST, the membranes were incubated with secondary antibodies conjugated with horseradish peroxidase at room temperature for 1h. After three washes with 0.05% TBST, immunodetection was performed with chemiluminescence as described above.

### RNA isolation

Cell pellets were homogenized in 1 ml of TRIzol reagent (Invitrogen, 15596026) and incubated at room temperature for 5 min. After incubation, 200 µl of chloroform was added, and samples were vortexed, incubated briefly at room temperature, and spun at 12000xg for 15 min at 4°C. The resultant aqueous phase was transferred to a new tube and mixed with an equal volume of 100% ethanol, then transferred to an RNeasy MinElute spin column (Qiagen, 74104). RNA was extracted on the column according to the manufacturer’s instructions. The quality of the eluted RNA was determined by measuring 28S/18S ribosomal ratios and RNA integrity number (RIN) using an Agilent Bioanalyzer.

### Real-time quantitative PCR

Reverse transcription of 1µg of total RNA was performed with random hexamers and SuperScript III reverse transcriptase (ThermoFisher #12574026) in a total volume of 20µl according to the manufacturer’s instructions. Reaction mixtures were incubated in a thermocycler at 25°C for 10 min then 50°C for 50 min before stopping the reaction at 85°C for 5 min. cDNA was then diluted 1:5 with nuclease-free water. Real-time quantitative PCR (qPCR) assays were performed in a total reaction volume of 20ul consisting of 4µl of diluted cDNA, 0.4µl of 10µM forward/reverse primer mix, 10µl of Power SYBR Green PCR Master Mix (Applied Biosystems 4367659) and 5.6µl nuclease-free water. Primer sequences used for qPCR are listed in the “Primers” subsection above. Reactions for each target gene were performed in duplicate in a 96 well plate loaded into an Applied Biosystems QuantStudio 3 Real-Time PCR System. Standard cycling conditions were used: Hold stage (x1): 50°C for 2 min, 95°C for 10 min; PCR stage (x40): 95°C for 15 sec, 60 °C for 1 min. Melt curve stage conditions were: 95°C for 15 secs, 60°C for 1 min, 95°C for 15 secs. Relative fold change in transcript abundance was calculated using the delta-delta Ct method by normalizing target gene expression levels to beta-actin. Significant differences in gene expression were evaluated using an unpaired t-test.

### eCLIP

AGO2 eCLIP was performed as described previously (Van Nostrand et al., 2016). Briefly, purified germ cell nuclei from adult mouse testes were crosslinked with 254 nm UV light. Nuclei were lysed and sonicated in a Bioruptor sonicator (Diagenode) at 4C, 5 min, 30 sec on/30 sec off. The nuclear lysate was then digested with RNase I (AM2295, Thermo) and Turbo DNase (AM2239, Thermo), and immunoprecipitated using anti-AGO2 antibody (04-642, Millipore). Next, the protein-bound RNAs were ligated with 3’ end adapters. Protein-RNA complexes were subjected to gel electrophoresis and transferred to a membrane, and RNA was isolated by cutting the membrane at the expected size range. cDNA was generated by reverse transcription, DNA adapter was ligated to the 3’ end of the cDNA, and the library was amplified by PCR. Screening of subclones prior to high-throughput sequencing was performed for quality control.

### Chromatin immunoprecipitation

Cells were cross-linked for ChIP-seq with 1% formaldehyde at room temperature for 10 min. The formaldehyde was quenched with 2.5M glycine at room temperature for 10 minutes. Fixed cells were spun down at 6000xg for 3 minutes at 4°C. Cells were washed with cold PBS twice and resuspended in 100ul ChIP lysis buffer (1% SDS, 10mM EDTA, 50mM Tri-HCl at pH 8.1) before being snap frozen in liquid nitrogen. Antibody-bound Dynabeads (Invitrogen 00821318) were prepared by mixing 10 μl aliquots of beads with 100 μl block solution (0.5% BSA in PBS). Beads were washed twice with 150 μl of block solution and resuspended in 30ul block solution. 1 μl of AGO2 antibody (Millipore, 04-642) was added to each aliquot of beads and incubated for 8 hours rotating at 4°C.

Between 2×10^5^ to 5×10^6^ cells were used for ChIP-seq. Frozen cells were thawed on ice and ChIP dilution buffer (0.01% SDS, 1.1% Triton X-100, 1.2mM EDTA, 167mM NaCl, 16.7mM Tris-HCl at pH 8.1) was added to reach a total volume of 300 μl. Cells were split into 150 μl aliquots in 0.5 ml Eppendorf tubes and sonicated at 4°C for 30 cycles (30 seconds on/off) using a Bioruptor (Diagenode). Aliquots of the same samples were pooled and spun down at 12,000xg for 5 min. The supernatant was moved to a new 1.5 ml Eppendorf tube and 600 μl of dilution buffer and 100 μl protease inhibitor cocktail (Roche #11836153001) were added. 50 μl of each sample was set aside as input before an aliquot of anti-AGO2 antibody bound Dynabeads was added and left overnight rotating at 4°C. After overnight incubation, beads were washed twice with low-salt immune complex wash buffer (0.1% SDS, 1% Triton X-100, 2mM EDTA, 150mM NaCl, and 20mM Tris-HCL at pH 8.1), twice with LiCl wash buffer (0.25M LiCl, 1% NP40, 1% deoxycholate, 1mM EDTA, and 10mM Tris-HCl at pH 8.1), and twice with TE (1 mM EDTA and 10mM Tris-HCl at pH 8.0). Bound DNA was eluted twice with 125 μl elution buffer (0.2% SDS, 0.1M NaHCO3, and 5mM DTT in TE) at 65°C and crosslinking was reversed with an 8 hour incubation at 65°C. ChIP and input samples were incubated for 2 hours at 37°C with 0.2 mg/ml RNAse A (Millipore 70856-3), and 2 hours at 55°C with 0.1 mg/ml Proteinase K (NEB P8107S).

Samples were prepared for sequencing using a Zymo ChIP DNA Clean & Concentrator kit (Zymo Research #D5201) according to the manufacturer’s instructions. Columns were washed twice with 200 μl wash buffer and DNA was eluted twice into fresh Eppendorf tubes: first with 7 μl of elution buffer and then with 6ul elution buffer.

### RNase Treatment

To treat mESC nuclei, we used a protocol adapted from (Sigova et al., 2015). Approximately 1×10^6^ mouse ESCs were grown in culture as described above. Cells were dissociated with 1x trypsin-EDTA 0.25%, spun down at 1500 rpm for 3 minutes, and resuspended in PBS. Cells were then washed twice with cold PBS, then resuspended in 5ml cold hypotonic solution (20mM HEPES, 20% glycerol, 10mM NaCl, 1.5mM MgCl2, 0.2 mM EDTA, 10% Triton x-100, 0.5 mM DTT, 1mM protease inhibitor, in H_2_O). Cells were incubated on ice for 10 min before nuclei were spun down and supernatant removed. The nuclear pellet was resuspended in 250 μl of cold PBS and incubated at room temperature with 15 μl RNase A (Millipore Sigma 70856-3) and 10 μl RNase H (New England Biolabs M0297S). The nuclei were prepared for ChIP as described above.

### Mass spectrometry analysis for AGO2 protein interactors

Nuclear and cytoplasmic fractions from wild type round spermatids purified by STA-PUT were isolated as described above and incubated with anti-AGO2 antibody (Millipore, 04-642). A parallel sample was incubated with normal mouse IgG as a negative control. Immunoprecipitated samples were then separated by SDS-PAGE and the gel was stained with Coomassie blue. All bands except immunoglobulin heavy chain and light chain were excised from the gel and subjected to mass spectrometry protein identification at the Keck MS & Proteomics Resource at Yale University. Briefly, individual excised gel bands in 1.5 ml Eppendorf tubes were washed 4 times; first with 500 µL 60% acetonitrile containing 0.1% TFA and then with 5% acetic acid, then with 250 µL 50% H2O/50% acetonitrile followed by a 250 µL 50% CH3CN/ 50 mM NH4HCO3, and a final wash with 250 µL 50% CH3CN/10 mM NH4HCO3 prior to removal of wash and complete drying of gel pieces in a Speed Vac. 10 µL of a 0.1 mg/mL stock solution of trypsin (Promega Trypsin Gold MS grade) in 5 mM acetic acid was freshly diluted into a 140 µL solution of 10 mM NH4HCO3 to make the working digestion solution. 124 µL of the working digestion solution was added to the dried gel pieces (additional 10 mM NH4HCO3 may be added to ensure gel pieces are completely submerged in the digestion solution) and incubated at 37 °C for 18 hours. Sample was then stored at −20 °C until analysis. Digested sample was injected onto a Q-Exactive Plus (Thermo Fisher Scientific) LC MS/MS system equipped with a Waters nanoACQUITY™ UPLC system, and used a Waters Symmetry® C18 180 µm x 20 mm trap column and a 1.7 µm, 75 µm x 250 mm nanoACQUITY™ UPLC™ column (37°C) for peptide separation. Trapping was done at 5 µl/min, 99% Buffer A (100% water, 0.1% formic acid) for 3 min. Peptide separation was performed with a linear gradient over 140 minutes at a flow rate of 300 nL/min. The data were processed and protein identification was searched using Proteome Discoverer (v. 2.2, ThermoFisher Scientific, Waltham, MA) and Mascot search algorithm (v. 2.6, Matrix Science LLC, London, UK). Mascot search parameters included: fragment ion mass tolerance of 0.020 Da, parent ion tolerance of 10.0 ppm, strict trypsin fragments (enzyme cleavage after the C-terminus of Lysine or Arginine, but not if it is followed by Proline), variable modification of Phospho Ser, Thr, and Tyr, Oxidation of Met, and Propioamidation of Cys, deamidation of Asn and Gln, and Carbamidomethlyation of Cys. Scaffold (version 4.8.6, Proteome Software Inc., Portland, OR). A protein was considered identified when Mascot listed it as significant and more than 2 unique peptides matched the same protein. The Mascot significance score match is based on a MOWSE score and relies on multiple matches to more than one peptide from the same protein. The Mascot search results were exported to an .xml file using a False Discovery Rate of 1% or less for the protein ID.

### Quantitative mass spectrometry analysis of protein expression

Pachytene spermatocytes or round spermatids were isolated by STA-PUT as described above and pellets containing 1-3×10^6^ pachytene spermatocytes or 0.4-1×10^7^ round spermatids were washed in PBS with protease inhibitors before sonicating in 400µl of RIPA buffer and then processed for Label Free Quantitative mass spectrometry at the Keck MS & Proteomics Resource. Briefly, the protein suspension was centrifuged at 14K rpm for 10 min, and 150 uL of the supernatant was removed. Chloroform:methanol:water protein precipitation was performed, and dried protein pellet was resuspended in 100 uL 8M urea containing 400mM ABC, reduced with DTT alkylated with iodoacetamide, and dual enzymatic digestion with LysC and trypsin (carried out at 37 C for overnight and 6 hrs., respectively). Digestion was quenched with 0.1% formic acid during the de-salting step with C18 MacroSpin columns (The Nest Group). The effluents from the de-salting step were dried and re-dissolved in 5 µl 70% FA and 35 µl 0.1% TFA. An aliquot was taken and concentration measured via Nanodrop, and diluted to 0.05 µg/µl with 0.1% TFA. 1:10 dilution of 10X Pierce Retention Time Calibration Mixture (Cat#88321) was added to each sample prior to injecting on the UPLC Q-Exactive Plus mass spectrometer to check for retention time variability and instrument QC during LFQ data collection.

LFQ data dependent acquisition (DDA) was performed on a Thermo Scientific Q-Exactive plus, Q-Exactive HFX, or Orbitrap Fusion Mass spectrometers connected to a Waters nanoACQUITY UPLC system equipped with a Waters Symmetry® C18 180 μm × 20 mm trap column and a 1.7-μm, 75 μm × 250 mm nanoACQUITY UPLC column (35°C). 5 µl of each digests (in triplicates) at 0.05 µg/µl concentration was injected in block randomized order. To ensure a high level of identification and quantitation integrity, a resolution of 60,000 was utilized for MS and 15 MS/MS spectra was acquired per MS scan using HCD. All MS (Profile) and MS/MS (centroid) peaks were detected in the Orbitrap. Trapping was carried out for 3 min at 5 µl/min in 99% Buffer A (0.1% FA in water) and 1% Buffer B [(0.075% FA in acetonitrile (ACN)] prior to eluting with linear gradients that reached 30% B at 140 min, 40% B at 155 min, and 85% B at 160 min. Two blanks (1st 100% ACN, 2nd Buffer A) followed each injection to ensure against sample carry over.

The LC-MS/MS data was processed with Progenesis QI software (Nonlinear Dynamics, version 2.4) with protein identification carried out using in-house Mascot search engine (v2.6). The Progenesis QI software performs chromatographic/spectral alignment (one run is chosen as a reference for alignment of all other data files to), mass spectral peak picking and filtering (ion signal must satisfy the 3 times standard deviation of the noise), and quantitation of proteins and peptides. A normalization factor for each run was calculated to account for differences in sample load between injections as well as differences in ionization. The normalization factor was determined by calculating a quantitative abundance ratio between the reference run and the run being normalized, with the assumption being that most proteins/peptides are not changing in the experiment so the quantitative value should equal 1. The experimental design was set up to group multiple injections (technical and biological replicates) from each run into each comparison set. The algorithm then calculates the tabulated raw and normalized abundances, and ANOVA *p* values for each feature in the data set. The MS/MS spectra was exported as .mgf (Mascot generic files) for database searching. Mascot search algorithm was used for searching against the Swiss Protein database with taxonomy restricted to *Mus musculus*; and carbamidomethyl (Cys), oxidation of Met, Phospho (Ser, Thr, Tyr), and Acetylation (N-term and Lys), and deamidation (Asp, Arg) were entered as variable modifications. Two missed tryptic cleavages were allowed, precursor mass tolerance was set to 10ppm, and fragment mass tolerance was set to 0.02 Da. The significance threshold was set based on a False Discovery Rate (FDR) of 1%, and MASCOT peptide score of >95% confidence. The Mascot search results was exported as .xml files and then imported into the processed dataset in Progenesis QI software where peptides ID’ed were synched with the corresponding quantified features and their corresponding abundances. Protein abundances (requiring at least 2 unique peptides) were then calculated from the sum of all unique normalized peptide ions for a specific protein on each run.

### High-throughput DNA sequencing

RNA sample integrity was assessed by BioAnalyzer (Agilent Technologies), and samples with RIN >7 were used for paired-end sequence library construction using the KAPA mRNA HyperPrep Kit (Roche # 08098123702). For ChIP samples, DNA integrity and fragment size were confirmed on a BioAnalyzer. Approximately 5-10 ng of DNA was end-repaired, A-tailed, adapter-ligated, and PCR enriched (8-10 cycles) using the KAPA Hyper Library Preparation kit (KAPA Biosystems, #KK8504). eCLIP libraries were prepared as described above according to (Van Nostrand et al., 2016). All indexed libraries were quantitated by qPCR using the KAPA Library Quantification Kit (KAPA Biosystems #KK4854). Samples were sequenced on an Illumina NovaSeq using 100bp paired-end sequencing or an Illumina HiSeq 2500 instrument using 75 bp single-end reads. Sample de-multiplexing was performed using CASAVA 1.8.2 (Illumina).

### ChIP-seq data analysis

ChIP datasets were quality filtered with the fastq_quality_filter tool from FASTX-Toolkit using parameters –q 20 –p 80 (Hannon, 2010) and quality was assessed using FastQC (Andrews, 2010). ChIP-seq data was aligned to the mouse genome build mm10 using Bowtie2 with --end-to-end --fast parameters (Langmead and Salzberg, 2012). Peaks were called with MACS2 at q=0.2 for round spermatid, spermatogonia, and mESC datasets, and q= 0.05 for pachytene spermatocyte datasets (Zhang et al., 2008). Pachytene spermatocyte replicates were overlapped using the intersect tool from Bedtools (Quinlan and Hall, 2010). For other cell types, the replicate with highest quality was used for analysis. Shuffled controls were generated by randomly re-assigning peaks in the dataset to new genome coordinates and leaving peak size unchanged, using the shuffle function in BedTools (Quinlan and Hall, 2010).

### RNA-seq data analysis

Raw sequence files were filtered and quality was assessed using FASTX-toolkit and FASTQC as described above. Filtered paired-end reads were matched to yield a file of common reads which were then pseudoaligned to the mouse (mm10) transcriptome from Ensembl release 102 (Howe et al., 2021) and quantified using kallisto v0.45.0 (Bray et al., 2016). Resulting transcript counts were summed to gene level and DEGs were called using DESeq2 with batch correction (Love *et al*. 2014). Lowly expressed genes with < 10 TPM were filtered from the analysis. Genes with fold changes ≥1.5 and adjusted *P* values ≤0.05 were considered differentially expressed.

### Splicing analysis

RNA-seq libraries were sequenced to a depth of 100 million reads to allow splice variant analysis. Splicing was assessed using rMATS with default parameters (Shen et al., 2014). Fastq files were used for rMATS input. Differential splicing calls and downstream analysis were conducted using the maser package in R (Veiga, 2018).

### eCLIP data analysis

AGO2 IP and size-matched input control (SMInput) samples were demultiplexed as previously described using the script demux_paired_end.py (Van Nostrand et al., 2016). 3’ and 5’ adapters were trimmed using Cutadapt v1.18 (Martin, 2011) with default parameters. To eliminate artifacts from rRNA and other repetitive sequences, trimmed reads were then mapped to RepBase (mouse mm10 RepeatMasker, last modified time: 05-May-2017 06:32) using STAR software v.2.6.1d (Dobin et al., 2013). Unmapped reads recovered after the RepeatMasker step were then mapped to the mouse mm10 genome assembly with options --outFilterScoreMin 0 -- outFilterScoreMinOverLread 0.33 --outFilterMatchNminOverLread 0.33 --alignEndsType Local. The script barcode_collapse_pe.py (Van Nostrand et al., 2016) was used to remove PCR duplicates based on unique molecular identifiers (UMIs). Peaks were called using the tag2peak.pl script from the CLIP Tool Kit (CTK) software (Shah et al., 2017) with an adjusted p-value threshold of 0.01 and normalized to the mm10 background model. Each AGO2 eCLIP replicate was also normalized to the corresponding SMInput data using the Peak_input_normalization_wrapper.pl script from (Van Nostrand et al., 2016). CTK was also used to track cross-link-induced mutation sites (CIMS) and cross-link-induced truncation sites (CITS) with a resolution of ±50nt at an empirical false discovery rate (FDR) of 0.01. Specific types of mutations, including deletions, substitutions and insertions, were detected by analysis of unique CLIP tags (trimmed mapped reads).

### Metagene analysis

Metagene analysis was performed using deepTools version 3.5.1 (Ramírez et al., 2016). Read distributions were compiled around peak centers. Raw H3K9me3 ChIP-seq data from meiotic cells (pachytene spermatocytes) was obtained from GSE137742 (Liu et al., 2019) and processed as described above. Raw ATAC-seq data from meiotic cells (pachytene spermatocytes) was obtained from GSE102954 (Maezawa et al., 2018) and processed as described above for ChIP-seq data. Locations of mouse recombination hotspots were obtained from DMC1 immunoprecipitation and sequencing-based detection of ssDNA (SSDS) data from GSE75419 (Smagulova et al., 2016) and intersected with meiotic AGO2 peaks using the intersect function from Bedtools (Quinlan and Hall, 2010). Shuffled peaks were generated using the shuffle function from Bedtools.

### Motif analysis

To obtain motifs enriched in ChIP peaks, a library of known motifs was screened using the findMotifsGenome.pl script in HOMER with default parameters (Heinz et al., 2010).

### Gene ontology (GO) enrichment analysis

Gene ontology (GO) enrichment analysis was performed using PANTHER GO annotations (DOI: 10.5281/zenodo.4495804 Released: 2021-02-01) at the GO consortium website (http://www.geneontology.org) (Mi, *et al*. 2017).

### Statistical analysis

Comparison of continuous variables was assessed using an unpaired t-test with Welch’s correction for duplicate and triplicate samples. Statistics for genomics and proteomics data analysis are described above.

## Data and code availability

All sequencing data has been deposited at GEO under accession number GSE181264. The mass spectrometry proteomics data have been deposited to the ProteomeXchange Consortium via the PRIDE (Perez-Riverol et al., 2019) partner repository with the dataset identifier PXD027935.

## Supplemental Information

**Figure S1.** Characterization of *Ago2* cKO testes.

**Figure S2.** AGO2 immunoprecipitation and mass spectrometry (IP-MS) in post-meiotic cell nuclear and cytoplasmic fractions.

**Figure S3.** Characterization of nuclear AGO2-bound transcripts identified by eCLIP.

**Figure S4.** Altered splicing and nuclear transcript retention in *Ago2* cKO germ cells.

**Figure S5.** AGO2 ChIP-seq in male germ cells.

**Figure S6.** Sperm motility and morphology in *Ago2* cKO.

**Table S1.** Differentially expressed genes in *Ago2* cKO

**Table S2.** Differentially expressed proteins in *Ago2* cKO

**Table S3.** AGO2 protein interactors identified by IP-MS

**Table S4.** AGO2 nuclear RNA targets identified by eCLIP

**Table S5.** Differentially spliced transcripts in *Ago2* cKO

**Table S6.** ChIP data summary, peaks, and associated genes

