## Supplemental Figures for "A nuclear role for the Argonaute protein AGO2 in mammalian gametogenesis"

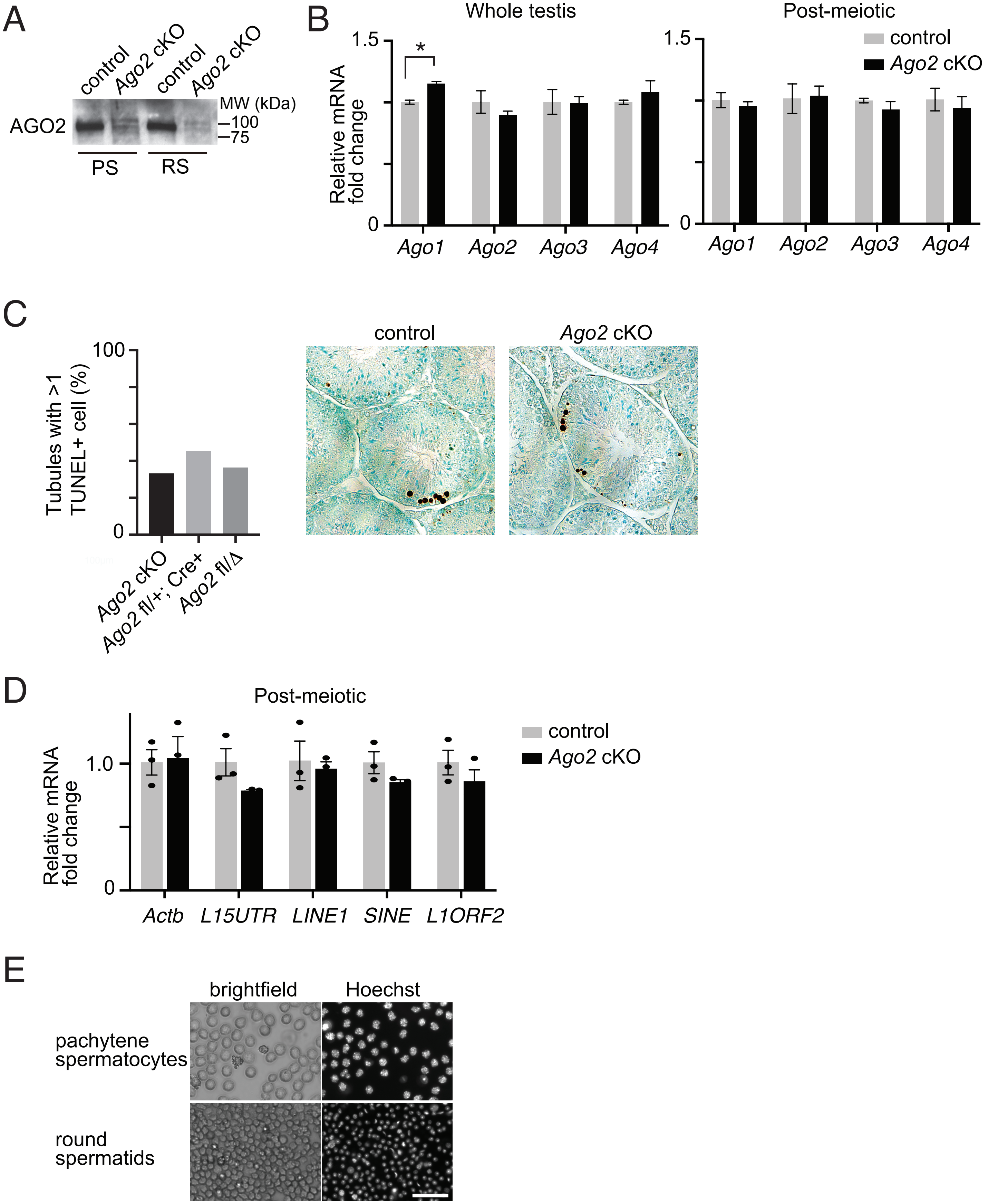


**Figure S1. Characterization of Ago2 cKO testes. A**, Long exposure of the Western blot shown in Figure 1A. **B**, RT-qPCR for all *Ago* family members in whole testes and post-meiotic cells of *Ago2* cKO and control males. **C**, Quantitation (left) and sample images (right) of TUNEL staining in *Ago2* cKO and littermate control (*Ago2* fl/+; Cre+ or *Ago2* fl/Δ) testes. **D**, RT-qPCR for LINE and SINE elements in *Ago2* cKO and control post-meiotic cells. **E**, Brightfield and Hoechst-stained images of meiotic (pachytene spermatocytes) and post-meiotic (round spermatid) cell populations isolated by STA-PUT. *p<0.05, Welch's t-test; all other comparisons not significant.

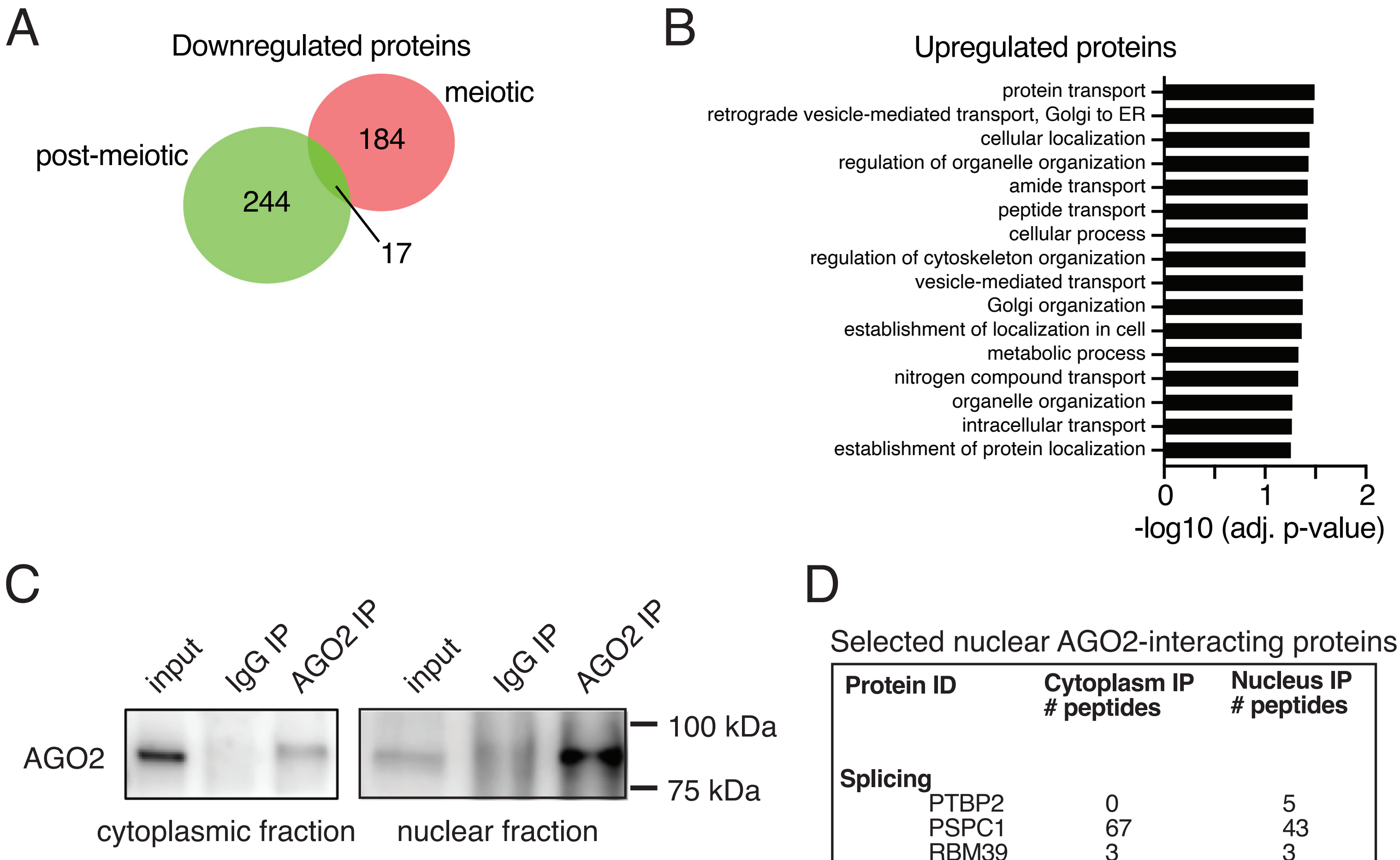


D

Selected nuclear AGO2-interacting proteins

| Protein ID | Cytoplasm IP # peptides | Nucleus IP # peptides |
|---|---|---|
| **Splicing** | | |
| PTBP2 | 0 | 5 |
| PSPC1 | 67 | 43 |
| RBM39 | 3 | 3 |
| SUGP2 | 3 | 22 |
| PRPF8 | 4 | 10 |
| HNRNPC | 2 | 11 |
| **Ribonucleoprotein nuclear export** | | |
| NUP210L | 16 | 25 |
| NUP160 | 3 | 14 |
| NUP98 | 7 | 15 |
| NUP133 | 3 | 9 |
| NXF1 | 5 | 23 |
| **Nucleosome positioning** | | |
| H1-2 | 3 | 7 |
| H1-3 | 2 | 4 |
| SMARCA5 | 0 | 3 |

**Figure S2. AGO2 immunoprecipitation and mass spectrometry (IP-MS) in post-meiotic cell nuclear and cytoplasmic fractions. A**, Overlap of proteins significantly downregulated in both meiotic and post-meiotic germ cells in Ago2 cKO males. **B**, Statistically enriched GO terms (FDR ≤ 0.05) associated with upregulated proteins in Ago2 cKO post-meiotic cells. **C**, Immunoprecipitation of AGO2 for IP-MS was validated in cytoplasmic and nuclear fractions of wild type round spermatids by Western blotting with anti-AGO2 antibody. 1% of total lysate before immunoprecipitation was used as input. Immunoprecipitation with mouse anti-IgG served as a negative control. **D**, Number of unique peptides detected following AGO2 IP-MS in wild type round spermatids for proteins associated with splicing, nuclear export, and nucleosome positioning.

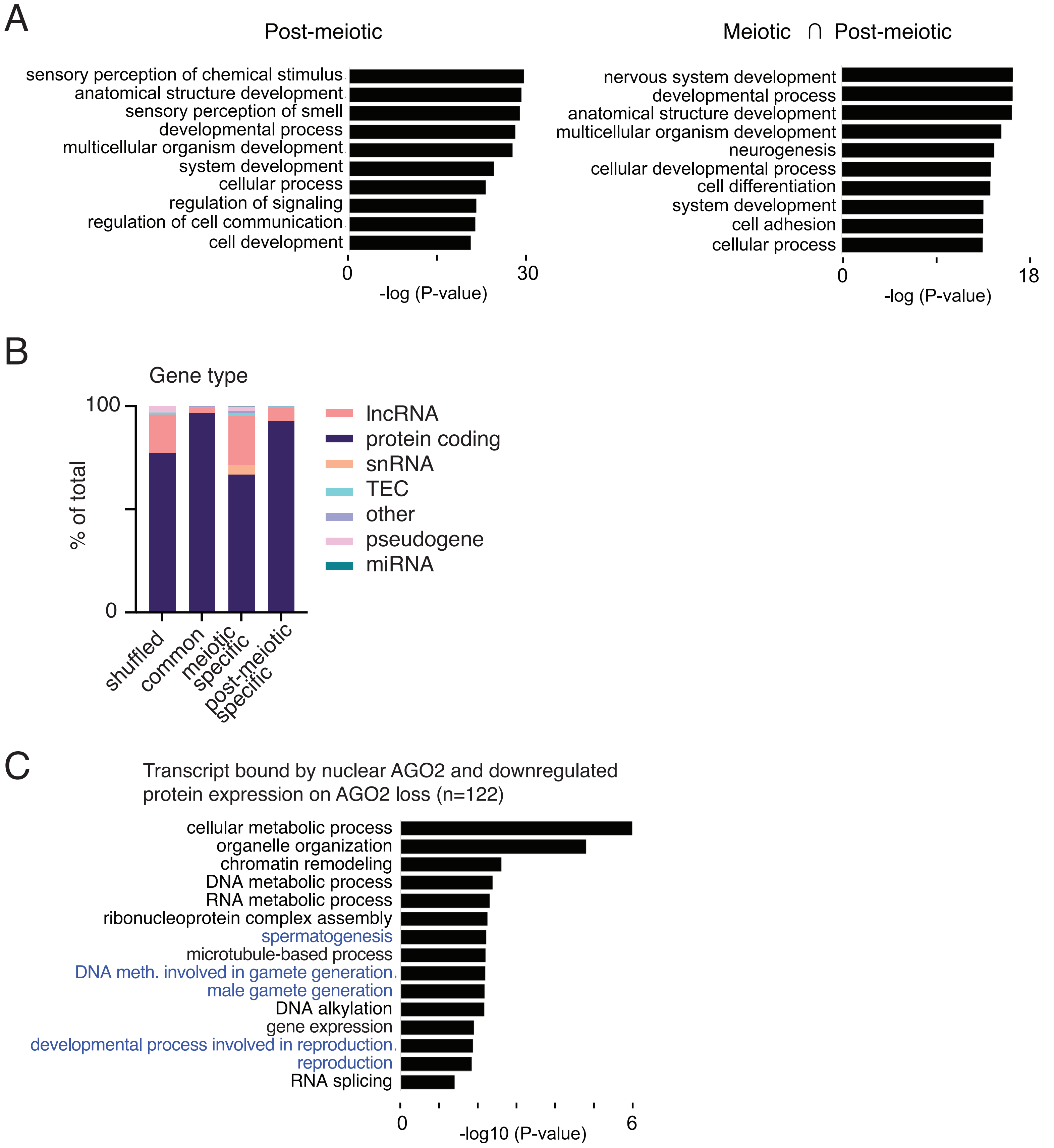


**Figure S3. Characterization of nuclear AGO2-bound transcripts identified by eCLIP. A**, GO analysis of post-meiotic consensus targets (left) and of targets bound in both meiotic and post-meiotic cells (right). **B**, Distribution of categories of bound transcript in both cell types (common), meiotic cells only, or post-meiotic cells only. The 'shuffled' control distribution was generated by randomly shuffling a representative CLIP peak set across all annotated transcripts. TEC, To be Experimentally Confirmed. **C**, Selected enriched GO terms for the sets of genes with downregulated protein expression and transcripts bound by AGO2 in meiotic or post-meiotic nuclei.

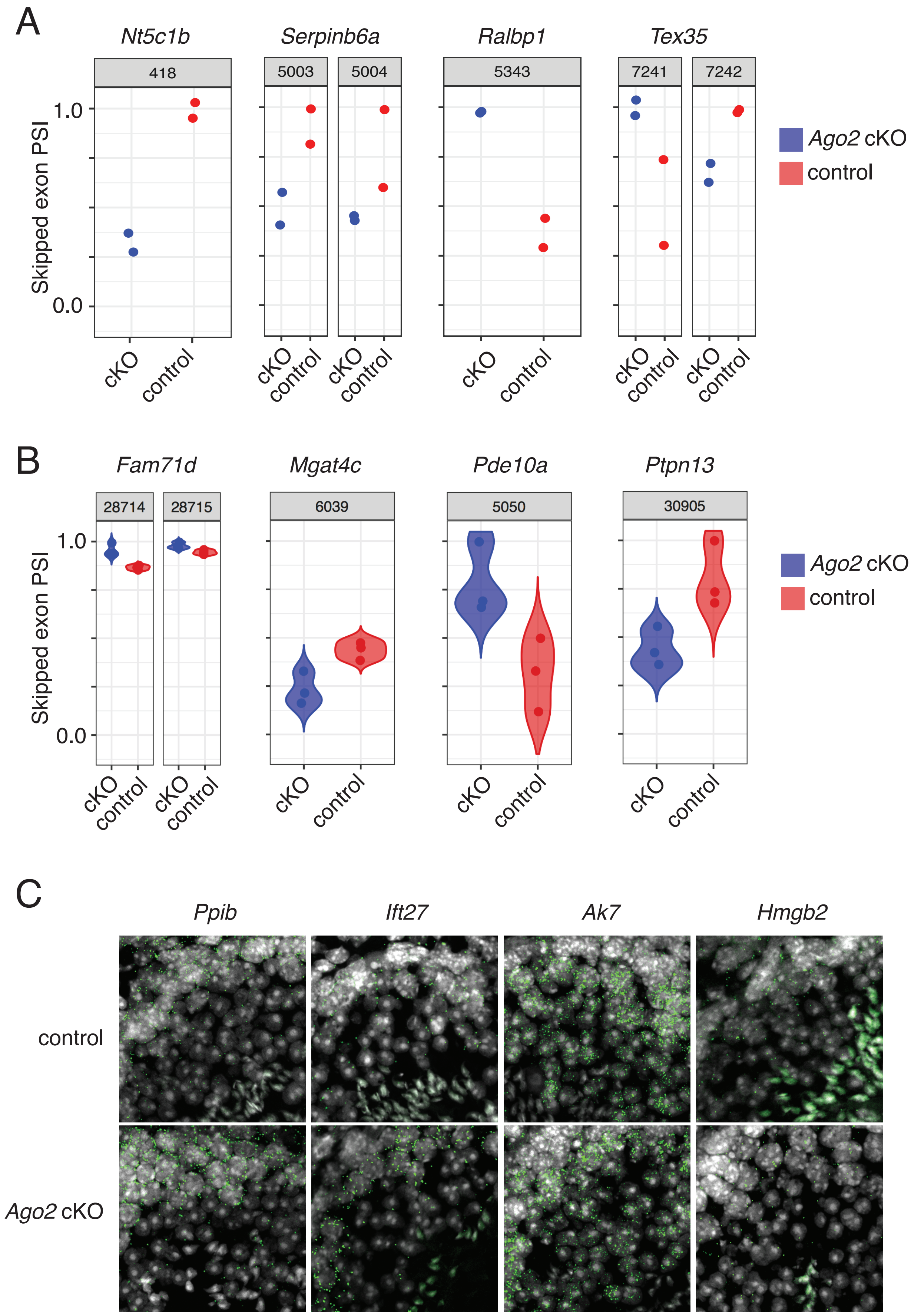


**Figure S4. Altered splicing and nuclear transcript retention in Ago2 cKO germ cells. A**, Examples of transcripts with differential exon skipping in control and Ago2 cKO meiotic cells. **B**, Examples of transcripts with differential exon skipping in control and Ago2 cKO post-meiotic cells. **C**, Sample RNAscope images in control and Ago2 cKO testes.

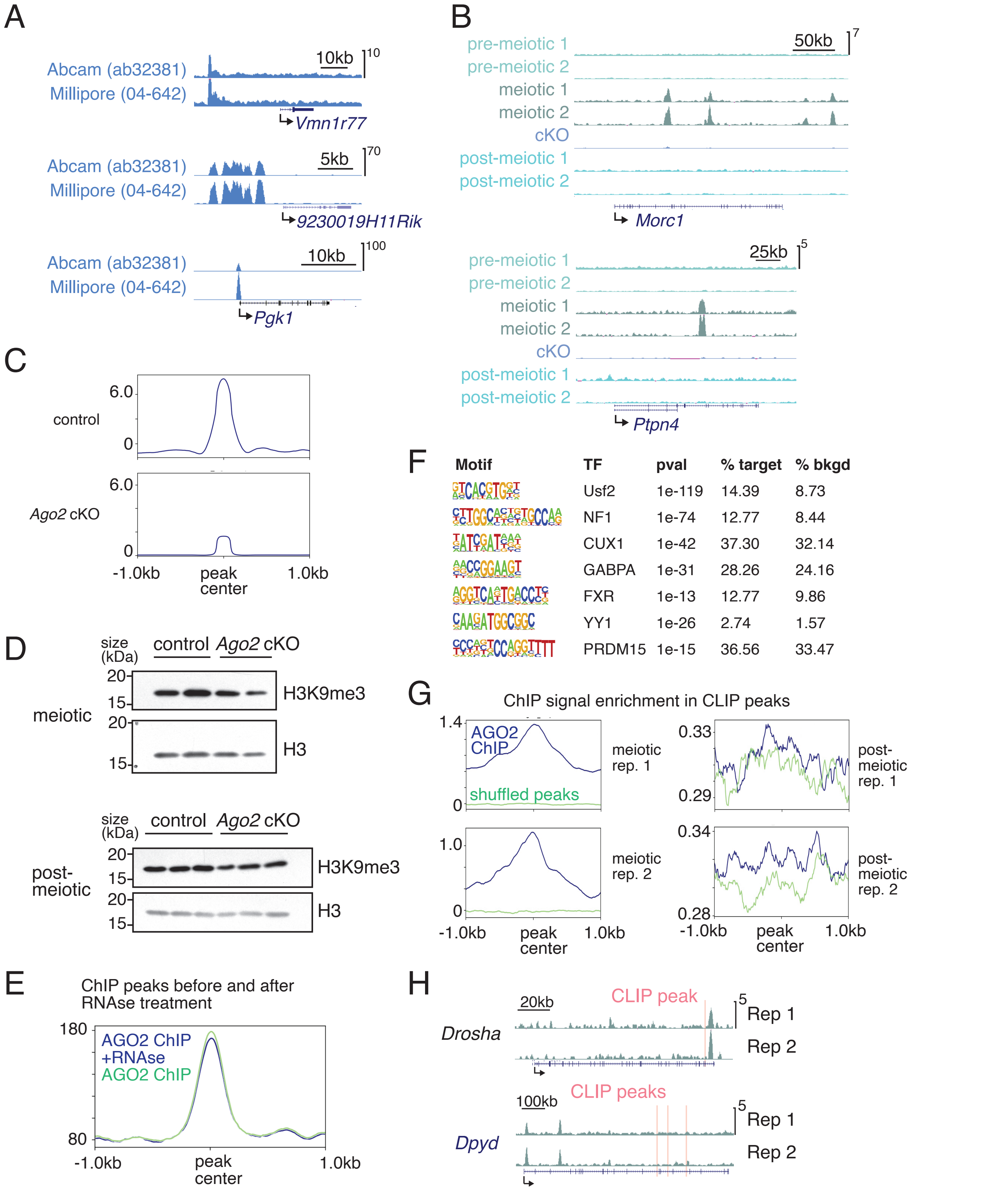


**Figure S5. AGO2 ChIP-seq in male germ cells. A**, Genome browser tracks from AGO2 ChIP seq in mESCs using two different antibodies. **B**, Additional representative genome browser tracks showing meiotic cell-specific AGO2 ChIP peaks in two biological replicates of pre-meiotic, meiotic, and post-meiotic wild type cells, and from Ago2 cKO mixed meiotic and post-meiotic cells. **C**, Metagene plots comparing ChIP signal in control and Ago2 cKO cells. **D**, Western blots showing global H3K9me3 levels in Ago2 cKO cells. **E**, Metagene plot comparing AGO2 ChIP signal in mESC nuclei treated with RNAse to an RNAse-negative control. **F**, Additional motifs enriched in meiotic cell AGO2 ChIP-seq peaks, where the corresponding transcription factors are expressed in meiotic male germ cells (see also **Figure 5F**). **G**, Separate metagenes for each biological replicate for AGO2 ChIP enrichment signal at eCLIP peaks(see also **Figure 5G**). **H**, Additional representative genome browser tracks for genes associated with both ChIP and CLIP peaks (see also **Figure 5H**).

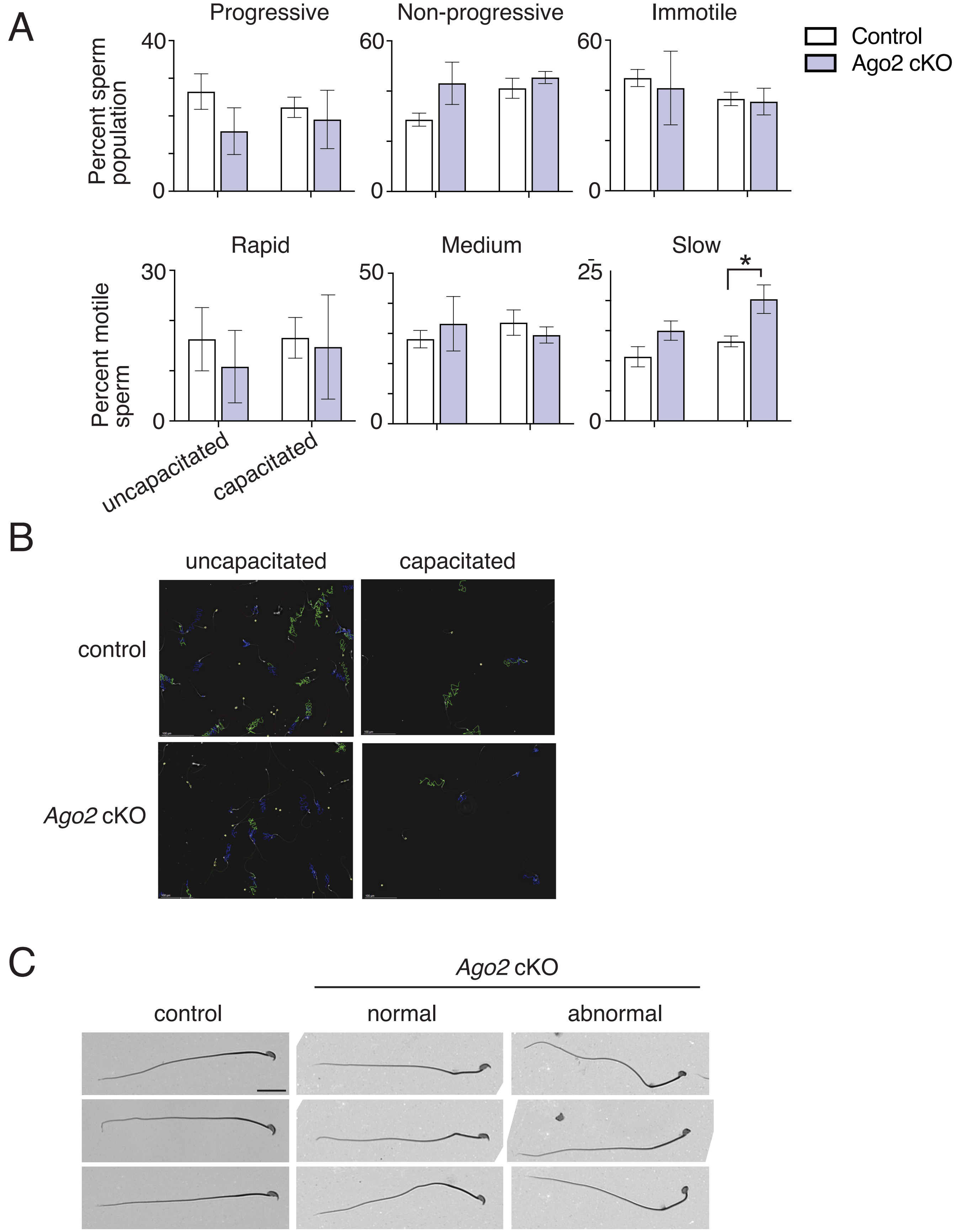


**Figure S6. Sperm motility and morphology in Ago2 cKO. A**, Results from computer-aided sperm analysis (CASA) showing motility parameters for control and Ago2 cKO sperm before and after capacitation. *p<0.05, Student's t-test. **B**, Representative images of control and Ago2 cKO sperm motility tracks. **C**, Brightfield images showing the full length of the spermatozoa with heads shown in Figure 6B. Scale bar, 20um.
